# Task engagement in immoral behavior altered post-task brain state

**DOI:** 10.1101/2025.10.12.681853

**Authors:** Eric Wang, Xinyi Julia Xu, Haiyan Wu

## Abstract

Studies have shown that the pattern of the brain in tasks can be effectively predicted from resting brain features. However, few studies explore how task could, in turn, influence resting state connectivity. This work attempts to understand this through the lens of immoral decisions. Using resting-state and task-state fMRI data collected before, during, and after an information-passing task involving dishonest choices with rewards, we employed the Hidden Markov Model (HMM) to capture the changes in the brain dynamics state. First, the HMM results identified 4 intrinsic brain states across both resting scans and task scans, including one reward network (state 1) and one control network (state 3). The reward network are associated with high dishonesty, while the control network are associated with low dishonesty. Importantly, task engagement in immoral behavior altered the post-task brain state. As participants engaged in persistent dishonesty, the network reconfigures itself to priorities the reward network and suppress the control network. Such suppression persists beyond the immediate task timeframe and leaves lingering signatures in the post-task resting scan. Contrary to the view of resting-state as a static baseline, our analysis showed the “non-resting” nature of post-task resting scans, where dishonesty could have an “after-effect” on neural dynamics. Together, these findings suggest that the ‘post-moral decision brain’ operates not as a static system but as a set of dynamically shifting states, whose adaptive trajectories may underlie the lasting effect of decision behavior in the prior task.

## 1 Introduction

Although the use of resting-state fMRI to predict subsequent task performance or neural activity is well-established, the converse question, how task engagement actively modifies intrinsic brain states, remains understudied. While task-based fMRI traditionally captures brain activity evoked by specific cognitive demands, the introduction of resting-state paradigms fundamentally transformed neuroimaging acquisition and analysis [6]. This shift enabled the identification of resting-state networks [52, 60], which have proven predictive of cognitive abilities [31, 10] and clinical phenotypes [17, 59]. Prevailing research, however, has largely focused on this unidirectional predictive relationship, overlooking the potential for tasks to exert a lasting influence on resting brain architecture. Growing evidence indicates that pre- and post-task resting states are neurophysiologically distinct, suggesting that task engagement may persistently alter intrinsic functional configurations [39, 54]. Task networks reshape resting-state networks by inducing subtle but functionally significant changes in brain connectivity and network architecture. Although resting-state networks (RSNs) provide a stable intrinsic framework, task engagement leads to selective reconfigurations, particularly in frontoparietal, cognitive control, and attention networks, which become more integrated and efficient during complex cognitive tasks [26]. These task-induced changes often involve a shift from more segregated resting-state modules to merged, task-specific modules that support increased cognitive demands, with enhanced connectivity between key networks such as frontoparietal, subcortical, and default mode networks [26, 13].

Importantly, evidence indicate that the efficiency of these network reconfigurations correlates with behavioral performance and cognitive abilities, such as reasoning accuracy and intelligence, suggesting that individuals with more optimized intrinsic networks require smaller, more efficient task-related updates [56]. Task-state functional connectivity changes, although modest in magnitude, significantly improve the prediction of cognitive task activations beyond resting-state connectivity, highlighting their functional relevance for dynamic cognitive processing [12]. Post-task resting states exhibit lasting modifications in network integration and hub roles, reflecting a task aftereffect that may enhance cognitive performance and differ between healthy and clinical populations, such as those with mild cognitive impairment [49]. Moreover, dynamic reconfiguration of frontal and frontoparietal networks—key regions implicated in executive control and moral cognition—supports the integration of affective and cognitive components necessary for moral decisions [7]. These findings suggest that efficient and selective brain network reconfigurations during and after moral decision tasks are essential for adaptive moral cognition and may serve as biomarkers of individual differences in moral behavior.

Moral decision task, involves complex cognitive processes that manage trade-offs between conflicting moral values and self-interest. Individuals show substantial variability in both behavioral responses and the neural processes when facing moral decisions [25]. Although some people adhere to universal ethical principles in their moral choices, others prioritize self-interest over moral standards [44, 23]. Indeed, previous research shows that the neural correlates of dishonest behavior and associated connectivity patterns also display considerable individual differences [25, 44, 23]. Specifically speaking, the neural correlates related to dishonesty include two important networks, that is, the cognitive control network and reward network. The former emerges as a pivotal cognitive mechanism in dishonest behavior, necessitating precise recall and increased cognitive resources to generate conflicting responses [64]. The latter acts as a major incentive in such dishonest behavior, the recruitment of the reward network often relates to increased dishonesty [36]. Together, the balance between the cognitive network and the reward network shapes the dynamic moral brain, which in turn influences the decision to put self-interest first or adhere to the moral norms. Indeed, individuals are shown to be incentivised by monetary gains to engage in dishonest behaviors [58, 21, 22, 84], and the interplay between moral norms and reward-driven motives can lead to behavioral inconsistencies [40, 58, 23]. Importantly, research also shows that instances of dishonesty may induce feelings of anxiety or guilt, leading to evident “after-effects” that impact subsequent behaviors and neural activity [33, 34]. One possibility we propose is that these aftereffects can persist beyond the immediate task time frame, leaving lingering signatures in functional connectivity and the brain network. It means that the post-task state “echoes” of the just-completed moral task, with a similar brain state pattern. Therefore, a post-task resting scan can act as a transition window, making it possible to capture these echoes of the just-completed task and to trace how transient, experience-dependent changes may reflect adaptations induced by the task, rather than a simple return to baseline.

Currently, relatively few studies simultaneously link (1) brain dynamics during an immoral decision task, (2) persistent after-effects of that task on the resting state of the brain, and (3) how these dynamic changes relate to behavioral tendencies such as immoral behavior. The present study aims to fill this gap by examining the full temporal arc of the moral brain by collecting pre-task resting scan as the cognitive baseline, task-based scan as task engagement, and post-task as brain adaptation, thereby characterizing how dynamics of brain reconfiguration. Recent work has demonstrated that brain activity is highly dynamic and can be clustered into discrete and functionally meaningful “brain states” [73, 42, 62]. The Hidden Markov Model (HMM) is one of the promising techniques to extract brain states from both task and resting fMRI sessions [73]. It assumes that the fMRI time series is generated by some hidden states in which the transitions are modeled as a Markov chain. It serves as a valuable tool to overcome the limitations that fail to capture time-varying changes in brain activity [74]. In recent years, HMM has been applied in various modalities, such as Eye-tracking [75], EEG [51], MEG [4], and fMRI [73]. In addition, HMM also demonstrates its capability in a diverse set of cognitive abilities, including attention [62], aging [45], schizophrenia [32], non-REM sleep [66], and more. These results demonstrate that HMM can exploit temporal information and have the potential to capture network reconfigurations before, during, and after motivated dishonesty.

To systematically investigate this directional interplay, specifically, how task networks reshape resting networks, we work on a dataset on motivated dishonest decision-making with additional pre- and post-task resting-state sessions. This longitudinal within-subject design enables us to trace how task-induced modifications of intrinsic brain states emerge and how such neural reorganization relates to behavioral outcomes. We applied HMM to investigate how the brain’s dynamic states, particularly those involving cognitive control and reward processing, are altered by engagement in motivated dishonesty and how these alterations manifest in the post-task period (Figure 1B). In the moral decision task, we manipulated the reward weights of the options to encourage dishonesty, honesty, or neither (random). Then, we applied HMM to explore the dynamic reconfiguration of large-scale functional brain networks, focusing on how brain state dynamics change before, after, and during the task under the conditions that encourage dishonesty (i.e., motivated dishonesty). Here we propose the following hypothesis (Figure 1A). First, the brain reconfigure its whole-brain network according to persistent task engagement, and the task-related changes will induce a carry-over effect in the post-task rest. This will be supported by the HMM analysis to identify distinct brain states common to both rest and task (H1). Second, brain reconfiguration would be associated with behavior during the task. The control network state will be negatively correlated with resistance to dishonesty, while the reward network state will be positively correlated with motivated dishonesty. This will be supported by complementary drift diffusion modeling (DDM) results linking the reward weight to decision bias (H2).

**Figure 1.**
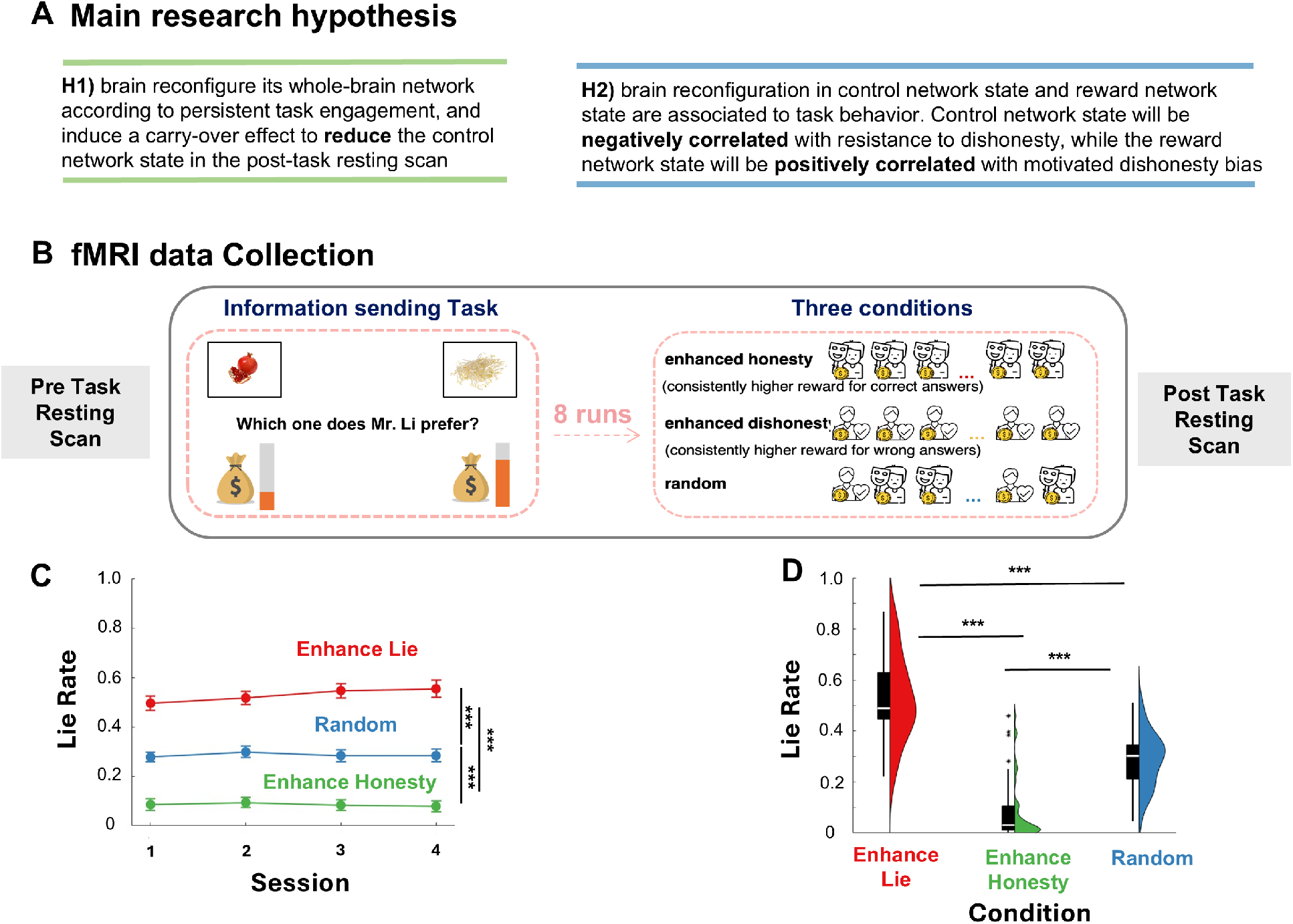
Experimental design with behavioral results. (A) Hypothesis of the current study (B) The fMRI data were collected during the whole experiment. There are 2 resting scans, each scan lasts 8 minutes, one collected before the information sending task, and one after the information sending task. In addition, there are also 4 scan sessions during the information sending task, where participants were asked to pass the preference information of Mr. Li to the next participant (the information receiver) with the consideration of reward units. We manipulated the reward unit differences between two items, thus yielding three conditions: enhanced dishonesty, enhanced honesty, and random. For details of the task, please see the supplementary materials. (C) Lie rate averaged separately for each of the four sessions and (D) across all sessions. The pair-wise comparisons are labeled in the figure, with “***” indicating *p* < 0.001. We found that all comparisons are significant, indicating our manipulations were successful. The result of the t-test corresponds to the F-tests in ANOVA.

## 2 Results

### 2.1 Behavioral performance

As indicated in Figure 1C, we found that our manipulation can evoke dishonest behavior. The lie rate shown here means the times that participants engage in dishonest behavior, where they report wrong answers. One-way analysis of variance reveals a statistically significant effect of the conditions on the lie rate. Importantly, this effect occurs both when we examined the data within each session separately (*F* = 60.33, *p <* 0.001 for session 1, *F* = 78.40, *p <* 0.001 for session 2, *F* = 82.81, *p <* 0.001 for session 3, *F* = 68.53, *p <* 0.001 for session 4) and when we collapsed across all sessions (*F* = 86.36, *p <* 0.001). Specifically, a Tukey HSD post hoc test was conducted, and the lie rate in the enhanced lie condition was significantly higher than in all other conditions, whereas the lie rate in the enhanced honesty condition was significantly lower than in all other conditions. The enhanced random condition consistently fell in between these two conditions. These results demonstrate that our manipulation reliably elicited the expected behavioral patterns in every session. In the following analyses, we focus on the lie rate in the enhanced lie condition, as it can be seen as a measure of motivated dishonesty.

#### 2.1.1 Decision process in the task

Our previous work suggests that the relative value of the history cumulative response (CR) has a significant effect on repeated moral decisions[81]. CR was used to quantify trial-by-trial consistency. It refers to repeating former choices, the time of choosing the same option across all repetitions[3, 57]. Thus the relative CR or consistency refers to CR of the wrong option minus CR of the correct option. We next used a Bayesian hierarchical drift-diffusion model (HDDM) framework suited for estimating trial-by-trial parametric modulations on latent decision processes[77] as our previous work [79]. We examined the extent to which evidence accumulation in the decision-making of IST was driven by relative reward and relative cumulative response (CR) between choice options. In the best model, the threshold was modulated by session number. This was to account for the significant reduction in reaction times along the session in IST (See Supplementary Figure. 3). The drift rate was weighted by relative reward and relative cumulative response (CR) in a trial-by-trial manner. The session-dependent initial bias term (z) was also added.

### 2.2 fMRI results

To capture the dynamics of brain states, we apply hidden markov models(HMM) to concatenate fMRI data. However, HMM faces great challenges of overfitting due to the high dimensionality of the data [2]. Therefore, based on previous studies [88], we train the HMM in the PCA space as shown in Figure 2. Specifically, the fMRI time series was reduced to retain 80% of the explained variance during training and were reprojected back onto the original Schaefer template space when training was complete [55]. Since estimating HMM requires specifying the number of hidden states, and there are no golden rules to determine this parameter [62, 45], we took an exploratory approach that not only considers the model fit but also the replicability of the HMM solution (see Supplementary Figure. 4-6 for model details). We found that the optimal states that are the most stable while still explaining the most were a 4-state solution. Similar to recent HMM studies [62, 9, 63], we also found a relatively low-dimensional state space.

**Figure 2.**
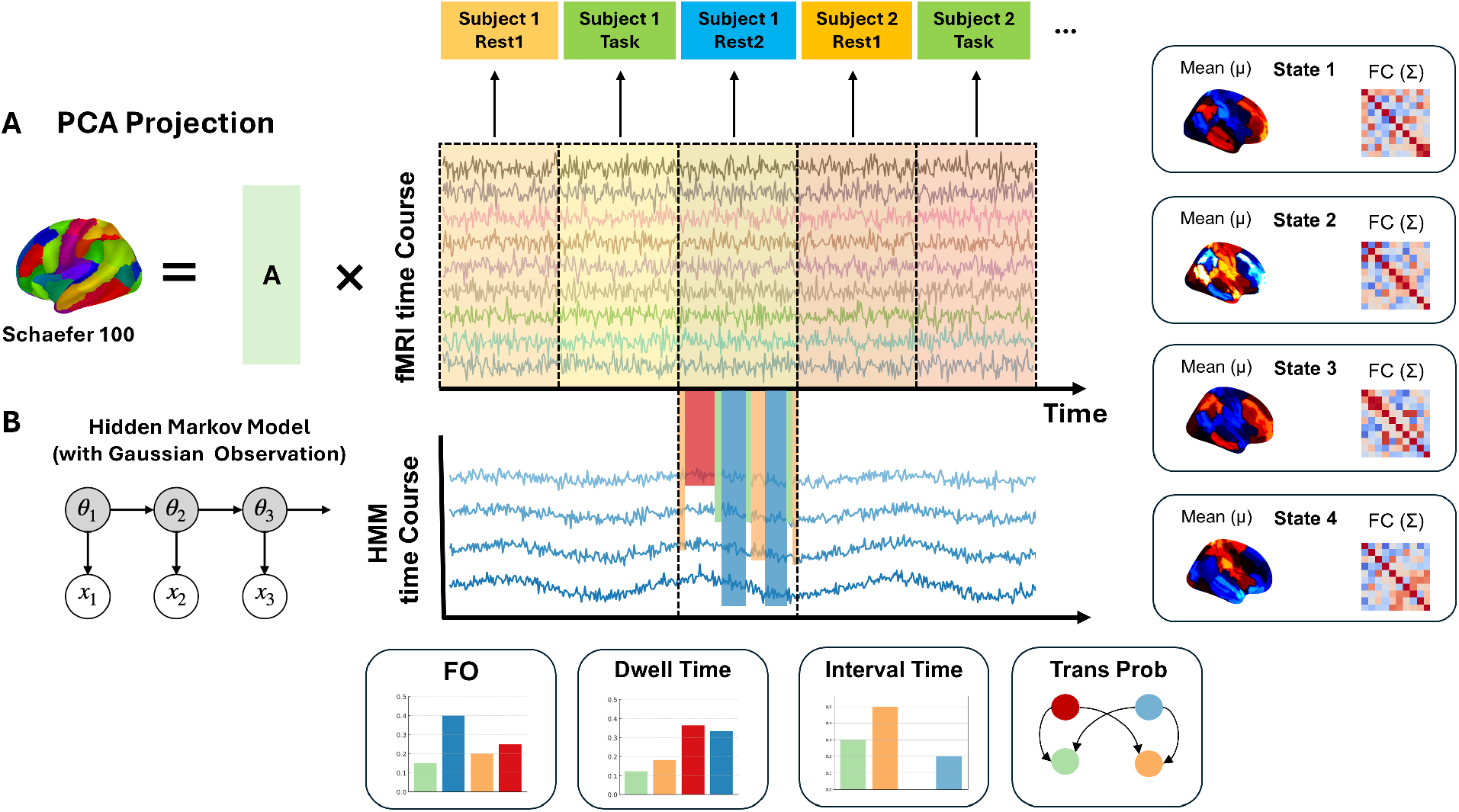
HMM pipeline on fMRI data. (A) The input is the ROI time courses extracted from the BOLD signals within each of the Schaefer 100 atlas for each participant. To address the model fitting challenges when estimating parameters, we train the HMM in PCA space by reducing its dimensionality. Specifically, 80% of the signal variance were kept. The time course were concatenated across participants across scan sessions. (B) We define HMM observation model to be a Gaussian Mixture model with 4 states. Each state is characterized by the mean and covariance matrix of a Gaussian distribution. In addition, it also yield a brain state time course where it specify which of the 4 states the current time point is current at. From there, we compute 4 dynamic states measures including the fractional occupancy, dwell time, interval time and transition probability.

#### 2.2.1 The profiles of 4 brain states occurring before, throughout and after the moral decision

Then, we investigated the relative weighting across the 100 atlas in each of the 4 states. To describe the functional relevance of the hidden states we performed the association mapping between the activation with 13 Neurosynth topics that contain a range of potential brain processes applicable to our task. Figure 3 displays the functional signatures of each brain state obtained from Neurosynth. We found 2 motor-related brain states, namely, state 2 and state 4. They were characterised by high correlation between sensorimotor and visual information. In addition, we also found state 1 was characterised by high correlation with keywords like reward and moral. On the other hand, state 3 was characterized by high correlation with keywords such as cognitive control, attention control, and conflict. Therefore, it seems that state 1 is a network that was a reward network. While state 3 was a control network. Therefore, we labeled state 1 as the “Reward Network”, state 2 as the “Motor Network”, state 3 as the “Control Network”, and state 4 as the “Motor Network 2”. For the mean activation map and functional connectivity of each latent state (see the supplementary Figure. 7). Since the training was completed on the concatenated data across scan sessions, the 4-states solution here can be viewed as the latent hidden space that organizes the dynamics before, throughout, and after the moral decision.

**Figure 3.**
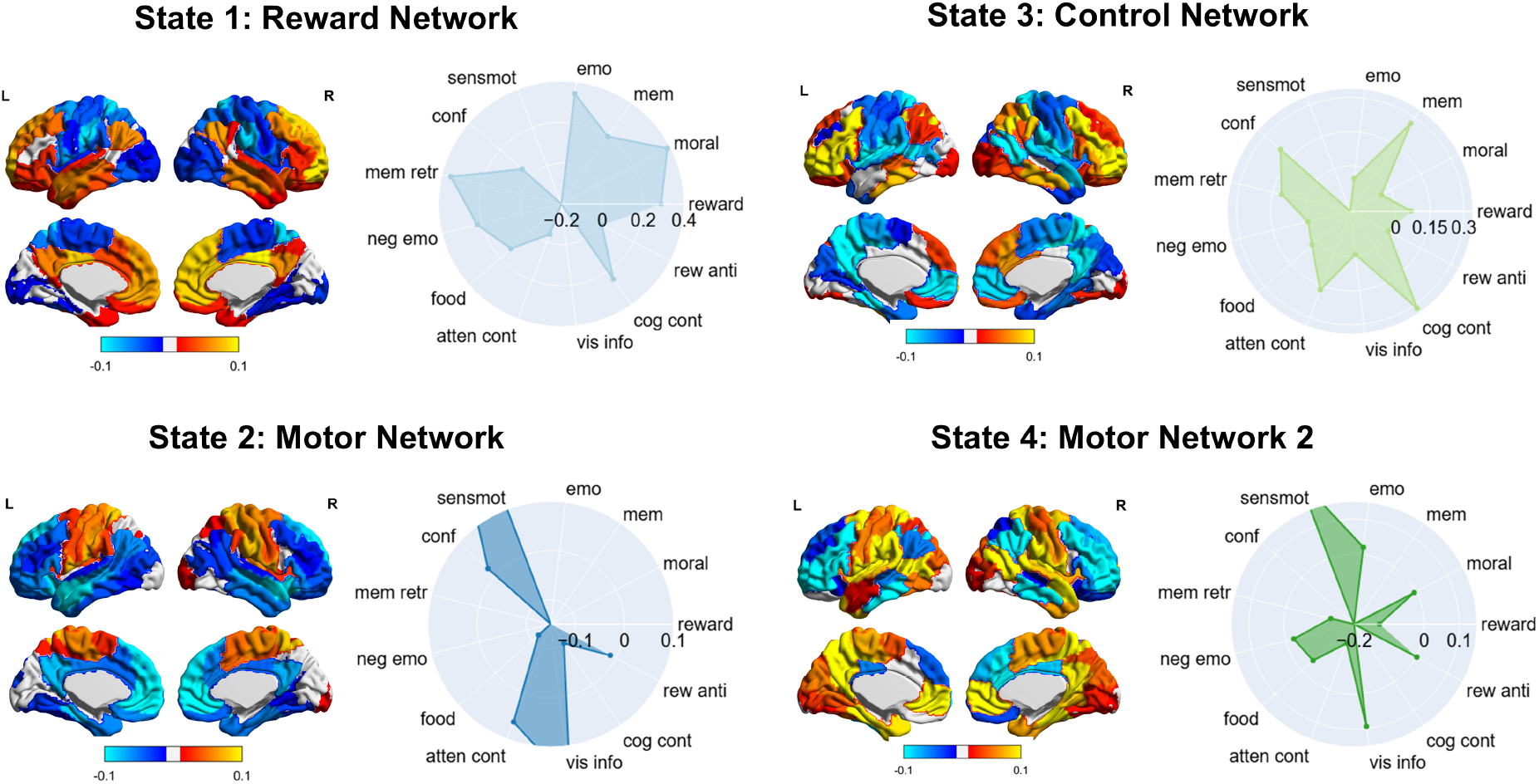
4 HMM states inferred by the HMM. The weighting on 100 atlas and their topic maps based on 13 Neurosynth keywords are shown. The score in topic maps represents the correlation between each brain state and Neurosynth topic. emo: emotion, mem: memory, moral: moral, reward: reward, rew anti: reward anticipation, cog cont: cognitive control, vis info: visual information, attend cont: attentional control, food: food, neg emo: negative emotion, mem retr: memory retrival, conf: conflict, sensmot: sensorimotor.

#### 2.2.2 Brain dynamically reconfigures its whole-brain network according to task

We then investigated changes in brain states during different scan sessions. Firstly, we compared the differences in HMM states in post-task rs-fMRI sessions and pre-task rs-fMRI sessions. Namely, we compared 4 HMM metrics: fractional occupancy(FO), dwell time, interval time and switching rate described in the method. As shown in Figure 4A, compared to the pre-task, we found that the post-task was characterized by significantly lower fractional occupancy in control network (*W* = 517, *p* < 0.05). No significant differences were found among other states after correcting for the false discovery rate. In addition, we also found that the overall switching rate between states drops significantly after the task (*W* = 485,*p* < 0.05) (Figure 4B). There were no significant results after FDR correction for dwell time and interval time. This shows that in general, the brain state after the task switches less between states, tends to be more stable, and shows a significant reduction in the time spent in the control state.

**Figure 4.**
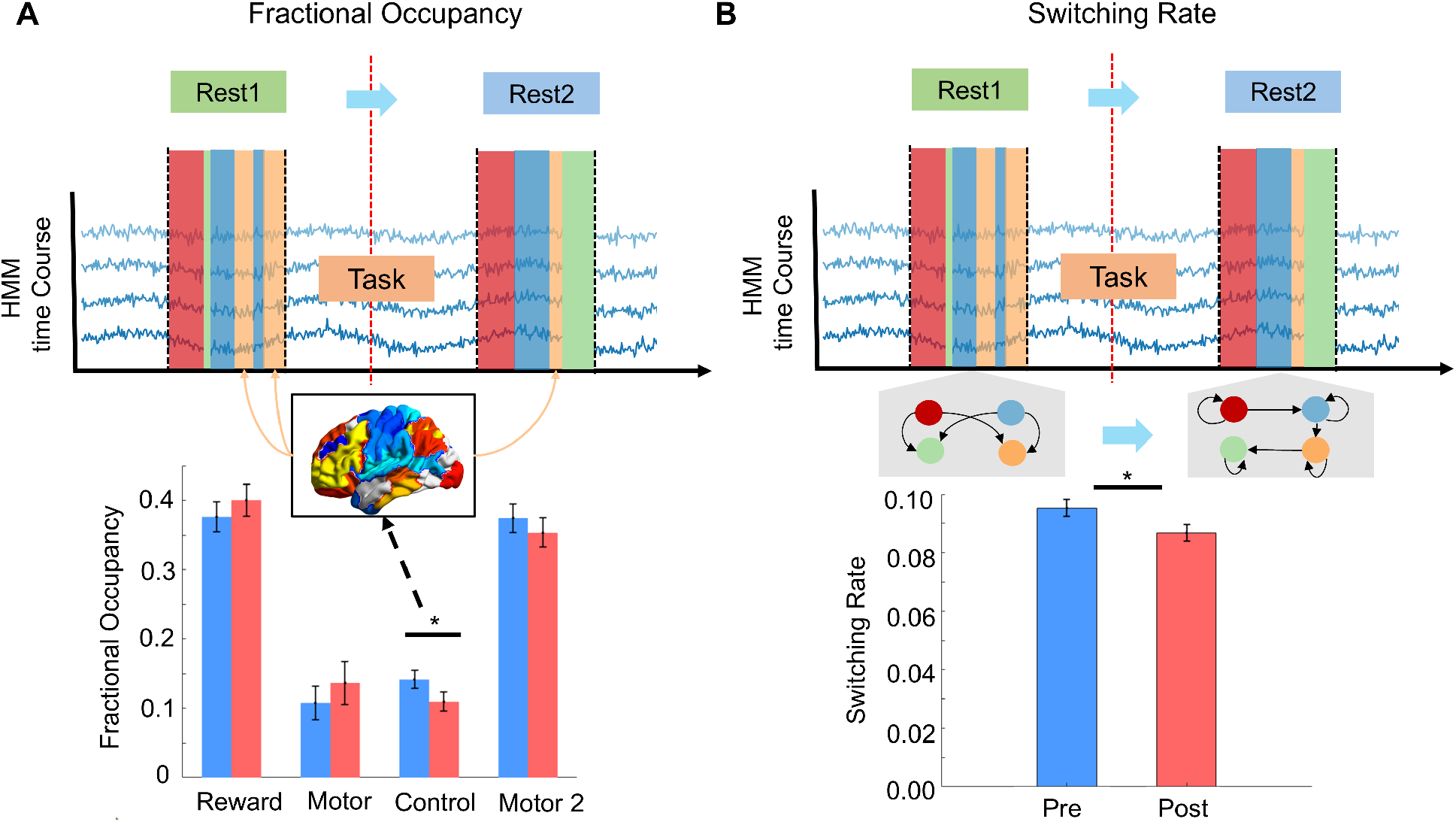
Comparison between pre-task rs-fMRI sessions and post-task rs-fMRI sessions. Asterisks denote statistical significance at (*p* < 0.05). Pre-task sessions are labeled in blue, whereas post-task sessions are labeled in red. (A) Top: Illustration of the reduction of the Fractional Occupancy, where the overall proportion of State 3 in orange decreased after the task. Bottom: The Fractional Occupancy of 4 States, where state 3 decreased significantly after the Task. (B) Top: Illustration of the reduction of the switching rate, where the between-state transition decreases after the task. Bottom: The switching rate of 4 states, where the switching Rate decreased significantly after the task

To further understand the neural dynamics, we compare the transition probability between different states in the pre-task, post-task, and during the task. As reported in Figure 5A, it is found that in pre-task and post-task HMM, the transitions are predominantly directed toward the reward network. In pre-task brain states, the highest transitions apart from self-transitions are from the motor network (0.0461), the control network (0.0701), and motor network 2 (0.0517) to the reward state. Similarly, in the post-task HMM, the transitions are predominantly directed towards the reward network, including transitions from the motor network (0.0569), the control network (0.0669), and motor network 2 (0.0484) to the reward network. By contrast, during the task, the transitions were more likely to converge to control network.

**Figure 5.**
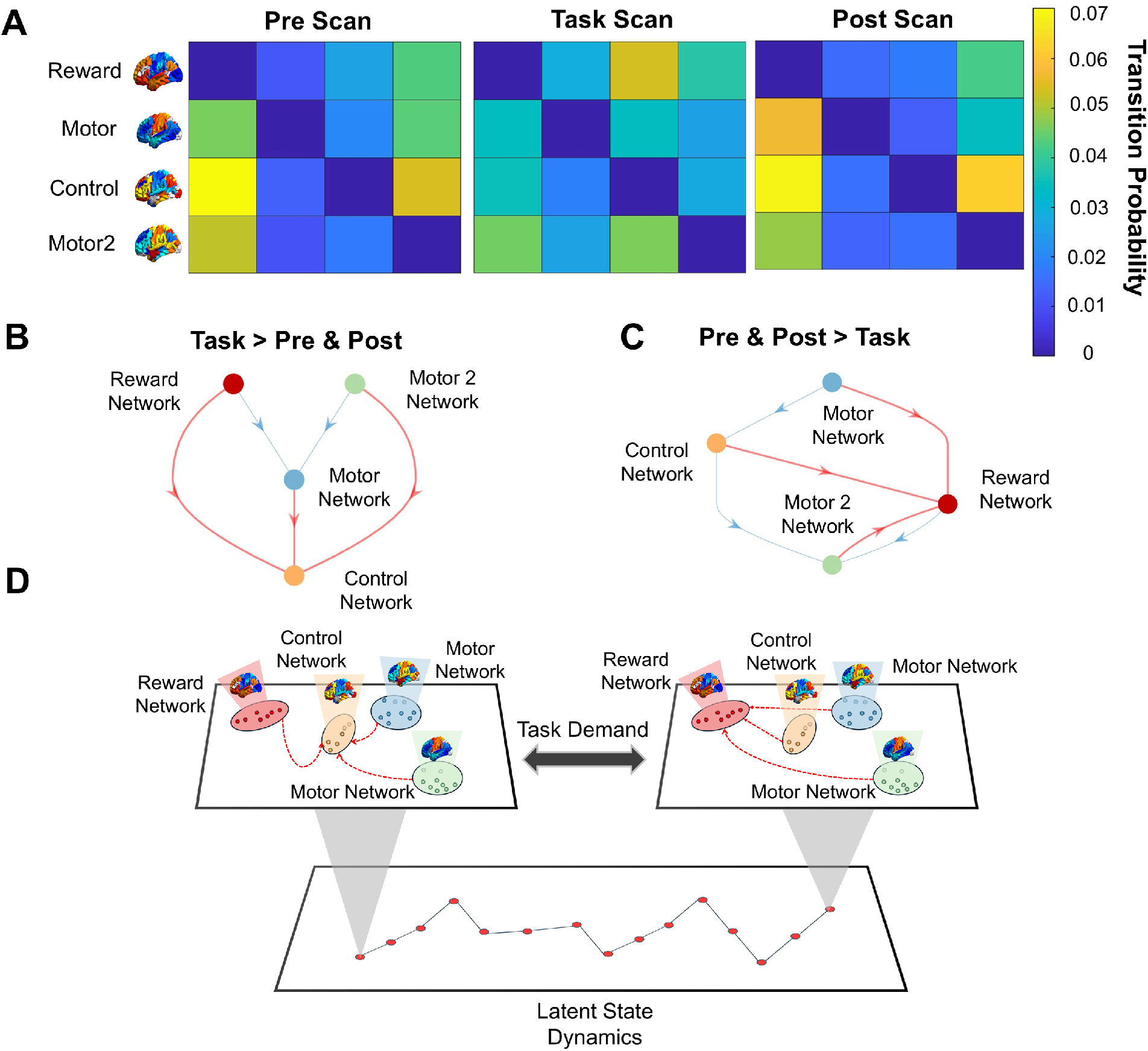
Comparison between pre-task rs-fMRI sessions, post-task rs fMRI sessions, and task fMRI sessions. Figures at top shows the transition probability between 4 states after removing self transition in (A) Pre-Task Scan session, Task Scan session and Post-Task Scan session. B) Statistically significant transitions detected using directional network-based statistics with significant level at (*p* < 0.05) characterized in task scan and (C) the transitions characterized in pre and post-task resting scans. (D) Figures in the bottom provides a summary of the latent dynamics that are modulated by task demand. In Task session, the control network acts as the hub node, where as in the pre and post-task scans, the reward network acts as the hub node.

For example, reward network tends to transit to the control network (0.0535) and the motor network 2 tends to transition to the control network (0.0465). Then, we applied directional network-based statistics (*p* < 0.05) to explore which of these transitions are statistically meaningful. The results in Figure 5B indicate that the transition probability in task HMM has the control network as a major “hub” node, where all other states are likely to transition towards the control network. Meanwhile, for the pre-task and post-task, the hub is replaced by the reward network Figure 5C. These transitions suggest that during the task, state transition favours the state of control state, while during the resting scans, the state transition favours the state of reward state. As illustrated in Figure 5D, this shows that the latent state dynamics vary across task and rest, with different configurations of the brain dynamics.

Finally, to more closely examine the dynamics of brain states during the task, we compared the fractional occupancy across different task scan sessions using repeated ANOVA. As shown in Figure 6A, the control network, compared to all other states, exhibited significantly higher fractional occupancy in sessions 1 and 2. However, as the task progressed with sustained motivational dishonesty, the fractional occupancy of the control network gradually decreased and eventually became non-significant in the later sessions. In contrast, the reward network gradually increased during the early stages of the task and then stabilized at a higher level in the later sessions. Figure 6B and Figure 6C offer a more close-up examination of the control state and reward state. As shown, the Tukey post-hoc test suggests there was a significant decrease from session 1 to session 2, and from session 2 to session 3. Similarly, the Tukey post-hoc test suggested there was a significant increase from session 1 to session 2. This indicates that, as the dishonest task unfolded, the engagement of the control network decreased, while the reward network became increasingly dominant (Figure 6D). Moreover, this observation is consistent with that the fractional occupancy of the control network decreased from pre-to post-task, suggesting that this reduction can be explained by the state dynamics induced during the task itself.

**Figure 6.**
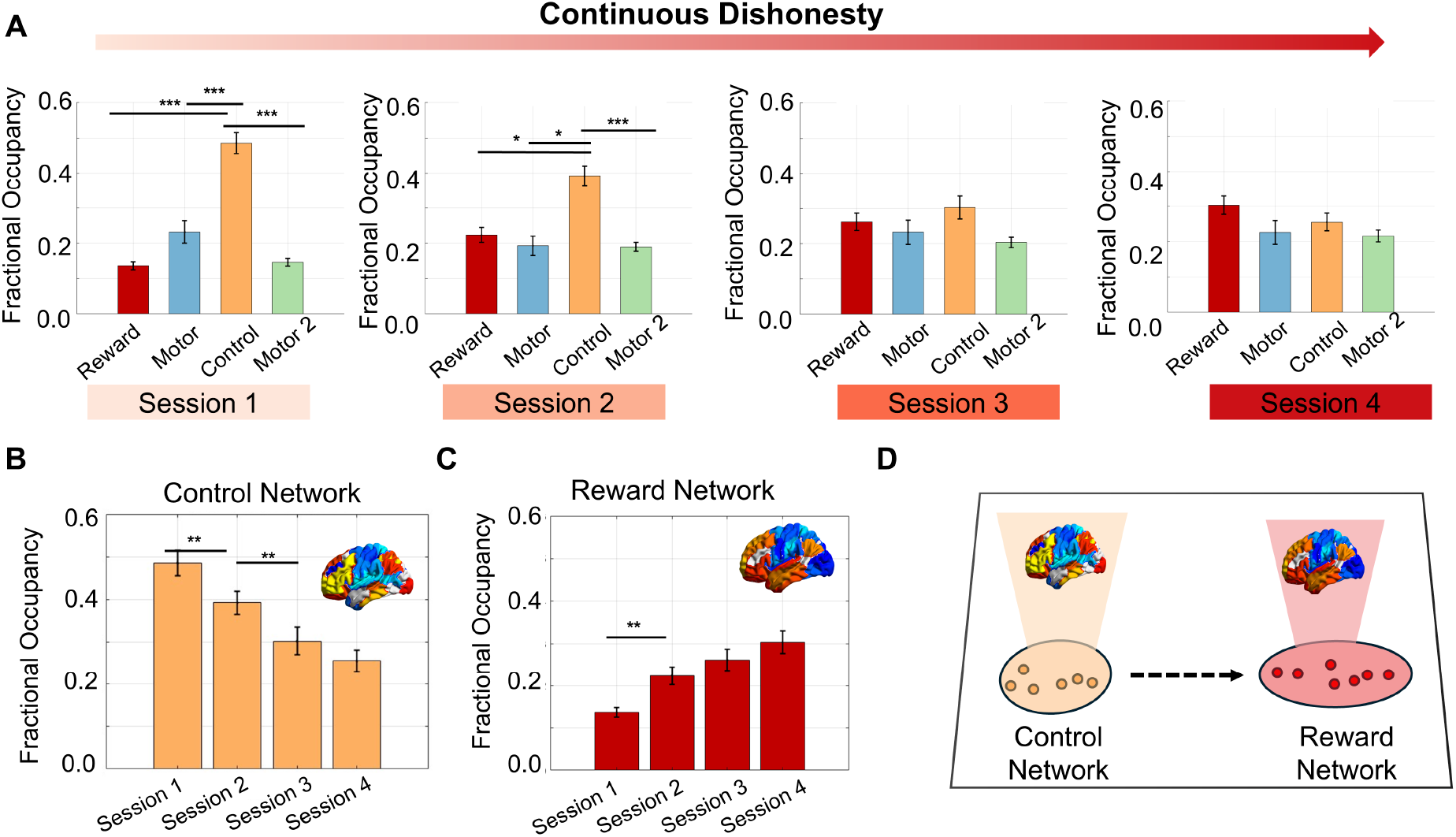
Comparison within the task fMRI sessions. (A) Figures at top shows the fractional occupancy of the 4 states in 4 different task sessions. (B) Figures at bottom shows the comparison for the control network and (C) reward network. Asterisks denote statistical significance at (*p* < 0.05, *p* < 0.01, *p* < 0.001). (D) Illustrates the overall tendency of reconfiguring brain dynamics towards reward network as dishonesty continue

So far, we have characterized the dynamic reorganization of brain states across pre-task, task, and posttask sessions, reflecting a redistribution of neural resources as motivational dishonesty unfolded. Collectively, these results indicate that the brain exhibits flexible reconfiguration of large-scale networks in accordance with task demands, and that such adaptive changes engender persistent modifications in subsequent resting-state dynamics.

### 2.3 Association between dishonesty behavior and Brain dynamics

Based on the observed impact of dishonesty on brain state dynamics, we next investigated whether these neural alterations reflect behavioral and neurodynamic correlates of motivated dishonesty. Specifically, we examined how motivated dishonesty—operationalized as the rate of lying under deception-conducive conditions—relates to four key metrics of brain dynamics: Fractional Occupancy(FO), Dwell Time, Transition Probability, and Interval Time. As mentioned above, we focus on the lie rate in the enhanced lie condition. This is because it directly incentivizes participants to lie in their answers for monetary gain. In doing so, it places participants in a moral conflict between pursuing self-interest and moral norms, thereby capturing the essence of motivated dishonesty.

In addition, we narrowed our analysis to states 1 and 3. Since the Neurosynth decoding revealed that state 2 and state 4 primarily reflected motor-related networks, which are less relevant to the moral decision process at the core of our study. By contrast, state 1 and state 3 were associated with higher-order cognitive networks, including reward processing and cognitive control. Given previous findings that both processes are critically implicated in motivated dishonesty [46], we therefore focused on these two states in the subsequent analyses.

#### 2.3.1 Relations between behavioral data and HMM dynamics

First, we investigate the relationship between the reward network and control network and the average lie rate across 4 task sessions. As indicated by the Figure 7A, we found that the dwell time of the reward network in session 1 positively correlates to the total averaged lie rate (*rho* = 0.45, *p* = 0.01). In other words, at the beginning of the dishonest task, spending more time in each visit to the reward network is related to a higher lie rate during the task. Moreover, we also found that the control network in post-task sessions is related to the total averaged lie rate. Specifically, Figure 7B shows that the fractional occupancy of control network negatively correlated to the averaged lie rate (*rho* = −0.37, *p* < 0.05), and Figure 7C shows that the interval time of control network positively correlated to the averaged lie rate (*rho* = 0.45, *p* = 0.01). This suggests that a greater proportion of the control network is associated with less lying during the task. Similarly, less frequent access to the control network after the task is associated with higher lie rate during the task.

**Figure 7.**
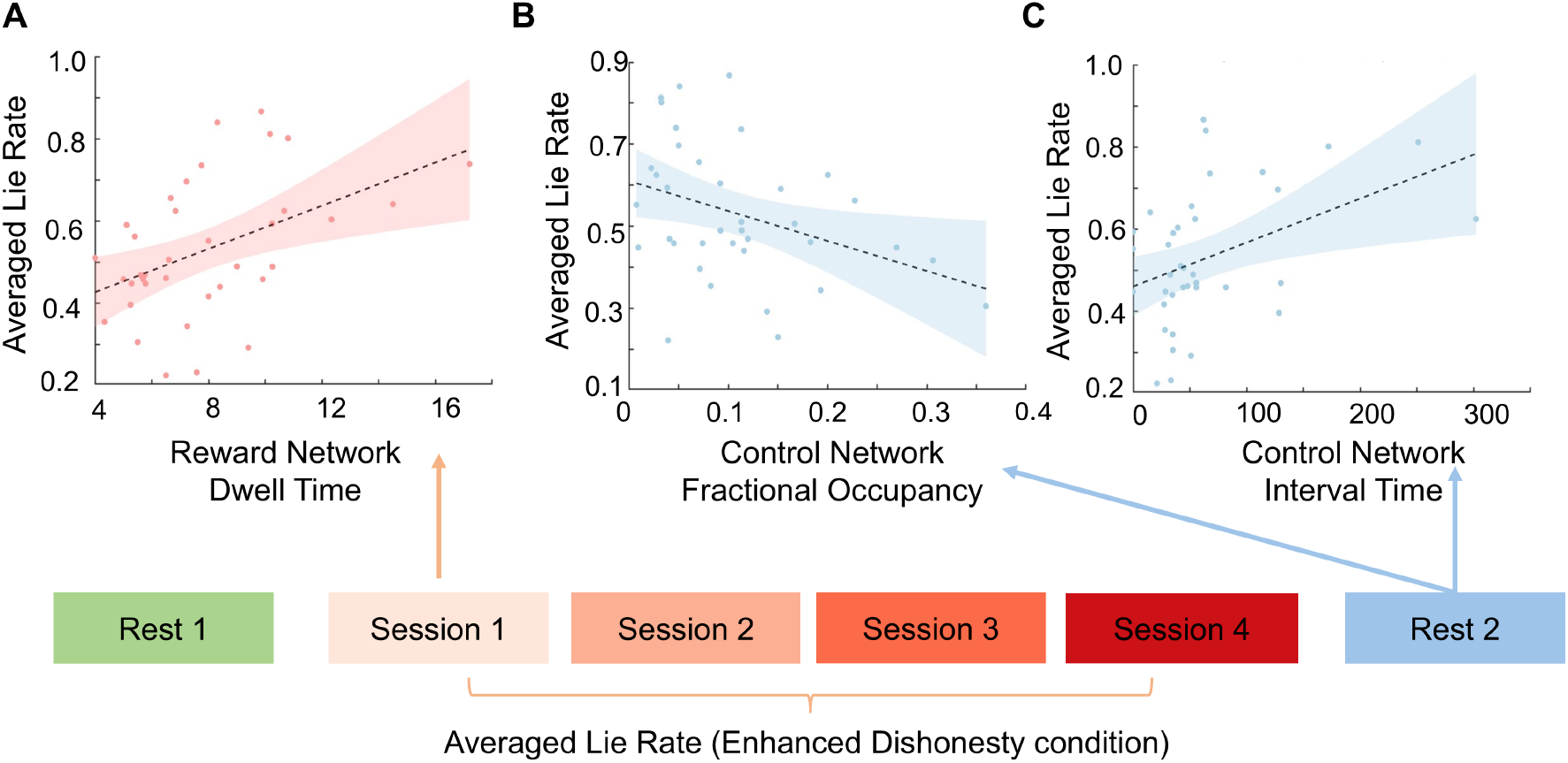
Reward Network and Control Network are related to lie rate averaged across sessions. (A) The relationship between Dwell Time of reward network from session 1 and total motivated lie rate (B) The relationship between Fractional Occupancy Time of Control Network and total motivated lie rate. (C) The relationship between Interval time of Control Network and and total motivated lie rate

We then extended this to the lie rate, session by session. We observed a significant relationship between 4 dynamic measures of Control Network in post-task with motivated dishonesty. In Figure 8A, we discovered that the fractional occupancy of Control Network negative correlated with the motivated dishonesty in both the session 1 (*rho* = −0.43, *p* < 0.05) and session 4 (*rho* = −0.44, *p* < 0.05) of the task. This demonstrates that, in the post-task resting state, individuals who spent more overall time in the Control Network were associated with a lower tendency to behave dishonestly during the task. In Figure 8B, we further found that the dwell time of Control Network in the post-task rest negatively correlated with motivated dishonesty in both session 1 (*rho* = −0.40, *p* < 0.05) and session 4 (*rho* = −0.40, *p* < 0.05). This indicates that individuals who stayed longer in the Control Network each time are associated with a lower tendency to lie during the task. In Figure 8C, inspired by previous work on HMM [87], we constructed a measure of cumulative transitions to Control Network by summing the transition probabilities from all other states into Control Network. We found that this cumulative transition measure in the post-task rest also negatively correlated with motivated dishonesty in session 1 (*rho* = −0.43, *p* < 0.05) and session 4 (*rho* = −0.41, *p* < 0.05), suggesting that the more frequently the brain state shifted into Control Network was associated with a lower tendency to behave dishonestly during the task. Finally, in Figure 8D, we observed a positive relationship between the interval time of Control Network in the post-task rest and motivated dishonesty in session 1 (*rho* = −0.41, *p* < 0.05) and session 4 (*rho* = −0.48, *p* < 0.01). This means that individuals who took longer to return to Control Network after leaving it were associated with a higher tendency to lie during the task.

**Figure 8.**
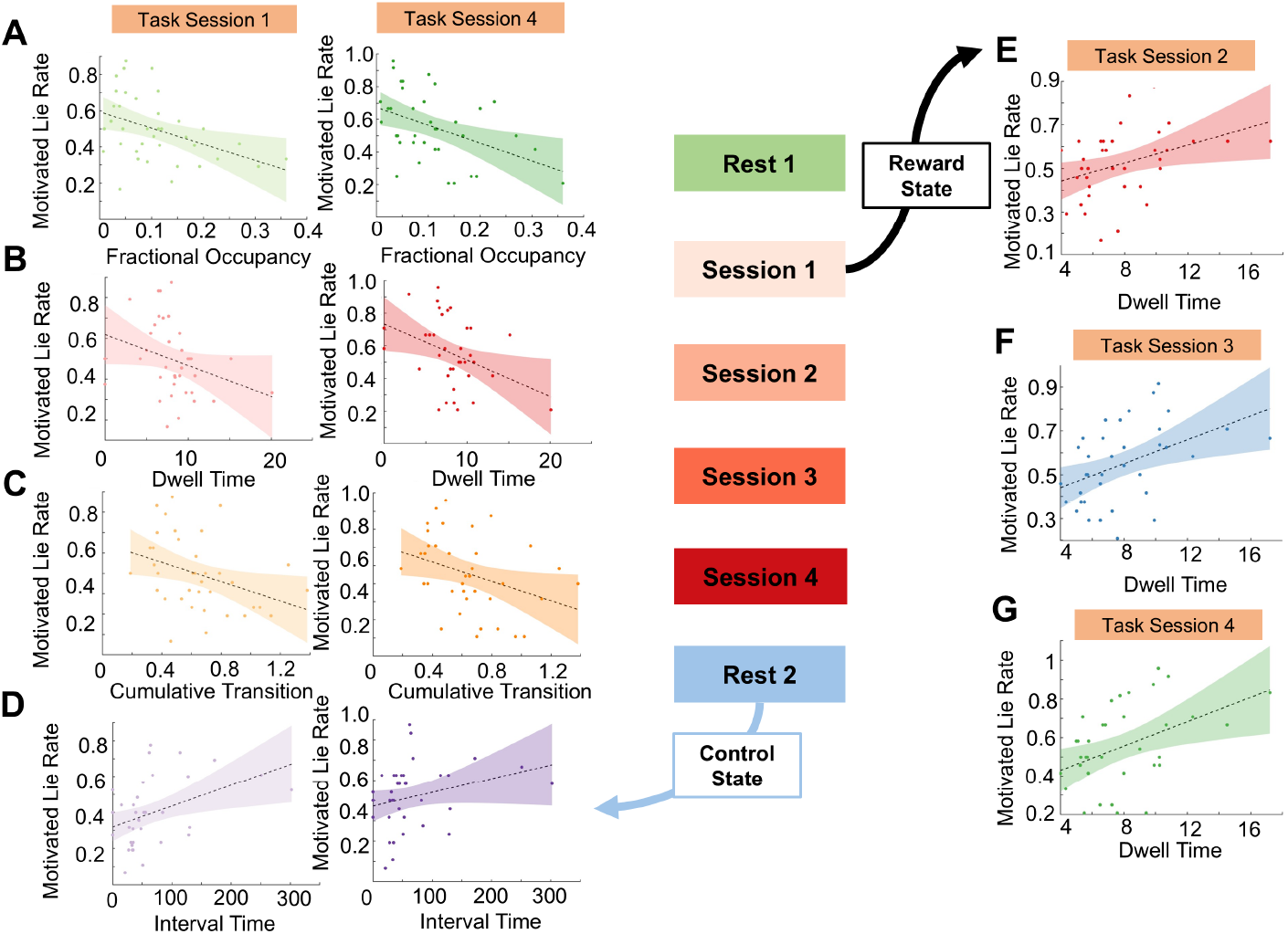
Post task control network activities are related to lie rate in both the start of the task and towards the end of the task. (A) The relationship between Fractional Occupancy of Control Network and motivated lie rate in session 1 and 4 of the task (B) The relationship between Dwell Time of Control Network and motivated lie rate in session 1 and 4 of the task. (C) The relationship between Cumulative Transition to Control Network and motivated lie rate in session 1 and 4 of the task (D) The relationship between Interval Time of Control Network and motivated lie rate in session 1 and 4 of the task. (E) In addition, the dwell time reward reward network in session 1 are related to motivated lie rate in session 2 (F) session 3, and (G) session 4

Importantly, none of these associations survived FDR correction in the pre-task resting state, and the effects were only present in the early and late stages of the task. Taken together, the findings suggest that after completing the task, greater engagement of the Control Network, which we interpreted as a control-related state, is associated with reduced motivated dishonesty.

For the Reward network, we observed a significant relationship in the dwell time in session 1 and the lie rate in the rest of the session. Specifically, as shown in the Figure 8E, the dwell time in session 1 positively correlated with the lie rate in session 2 (*rho = 0*.*43, p* < 0.05). Similarly, it also positively correlated with lie rate in session 3 (*rho = 0*.*50, p* < 0.01) and session 4 (*rho = 0*.*41, p* < 0.05) shown in Figure 8F and Figure 8G. The result here seems to suggest that the degree to which the reward network was activated when first encountering the task influences the lie rate in subsequent sessions.

In general, these results revealed that activating the reward network at the beginning of the task was associated with dishonesty in the subsequent task, while activating the control network after the task was associated with honesty in the task.

#### 2.3.2 Relations between DDM Results and HMM dynamics

To further understand the relationship between the brain state dynamics and behavior, as shown in Figure 9A, we utilized the drift diffusion model to explore the behavioral dynamics. The DDM parameters used are *V*_*Diff*_ and *Z*_*Ses*_. The *V*_*Diff*_ is the interaction term between the evidence cumulation rate (Drift rate) and the reward. The *Z*_*Ses*_ is the interaction term between the starting point (response bias) as the session unfolds. Here, *V*_*Diff*_ can be interpreted as the weight of the monetary reward, and *z*_*Ses*_ can be interpreted as how people change their bias for their decision as dishonesty continues.

**Figure 9.**
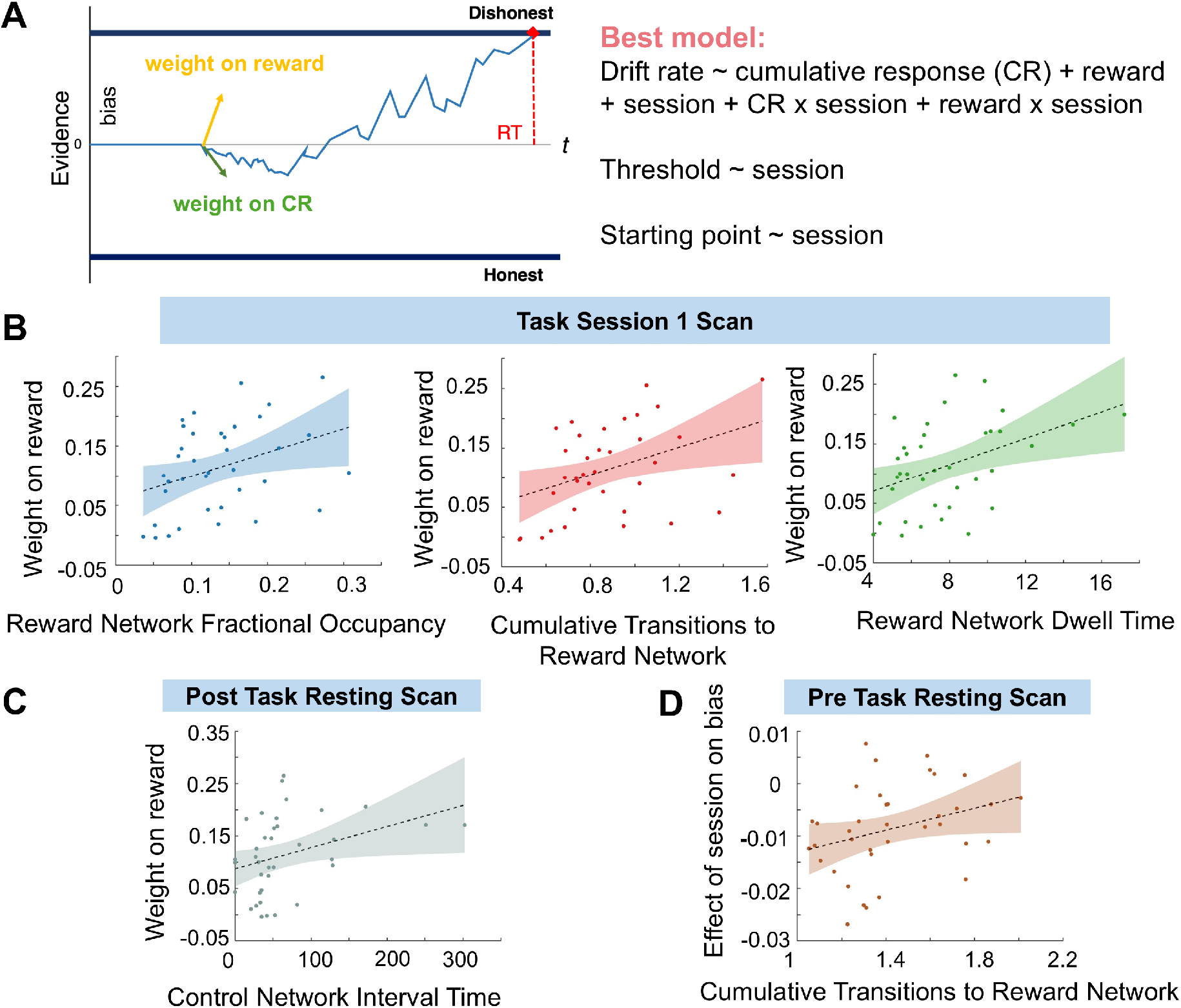
Results of Brain dynamics and behavioral dynamics. (A) Illustration of the DDM parameter used in the analysis. (B) The relationship between weight on reward and reward network dynamics in task session 1 scans (C). The relationship between weight on reward and control network interval time in the post-task resting scan (D). The relationship between the effect of the session on bias and the cumulative transitions to the reward network in the pre-task resting scan.

Figure 9B illustrates the relationship between the dynamic metrics of the reward network at the early stage of the task (session 1) and the weight on reward. We discovered that fractional occupancy (*rho = 0*.*42, p* < 0.05), cumulative transitions (*rho = 0*.*42, p* < 0.05), as well as dwell time (*rho = 0*.*43, p* < 0.05) of the reward network positively correlate with weight on reward. This demonstrates that individuals who spent more overall time in the reward network, or individuals who more frequently shifted into the reward network, or stayed longer in the reward network in each visit, will have more weighting on the monetary value in their decision. Moreover, we also found that in the post-task resting scan, the interval time of the control network positively correlated with weight on reward Figure 9C (*rho = 0*.*45, p* = 0.01). This suggests that individuals who place more weight on the reward value in their decisionmaking are associated with reduced visiting frequency of the control network in the post-task session. Finally, as shown in Figure 9D, we found that the cumulative transitions to reward network positively correlated with the effect of session on bias in pre-task resting scans (*rho = 0*.*37, p* < 0.05). This suggests that the overall tendency of transitioning into the reward network before the task begins is related to more biased, dishonest decisions. These results demonstrate that the brain dynamics of the reward network in the early stage of the task, both in the pre-task resting scan and session 1 of the task scan, are associated with a dishonest decision tendency and more reward-driven decisions. Moreover, we also extended these findings by highlighting the role of the control network. Specifically, the less frequently controlled network was visited after the task — that is, the longer the intervals between successive visits, the more strongly individuals tend to weight rewards in their decisions. In sum, by linking behavioral measures with neural state dynamics, our results suggest that reward network activity is associated with dishonest, reward-driven choices. In contrast, the control network is more closely related to honest decision-making.

## 3 Discussion

Task engagement induces subtle but functionally significant reconfigurations of resting-state brain networks, particularly for networks involved in the task. Dishonesty, a pervasive behavior in everyday life, has been extensively studied and identified engagement of cognitive control and reward brain network in it [40, 18, 68]. In contrast, the task-free or “resting” state has classically been interpreted as the brain’s intrinsic, baseline configuration. Here, we explored how task engagement would alter brain network at rest. Unlike the pre-task period, the post-task period provides a critical window into how a sustained experience, such as repeated dishonesty, induces persistent adaptations in the brain’s intrinsic dynamic repertoire. Based on a comprehensive fMRI data and HMM Figure 10A, this study systematically manipulated reward contingencies in different trials to examine how motivated dishonesty reshapes the dynamics of brain states. To our knowledge, this is the first time a study has compared the dynamic reconfiguration of brain states before, during, and after the task and linked these changes to neural and behavioral outcomes. We identified four promising preliminary findings. First, we find a reduction in the fractional occupancy (FO) of the control network after dishonesty, accompanied by a global decrease in brain state switching rates. Second, transitions during the dishonest task tend to converge on the control network, whereas transitions during resting states preferentially converge on the reward network.

**Figure 10.**
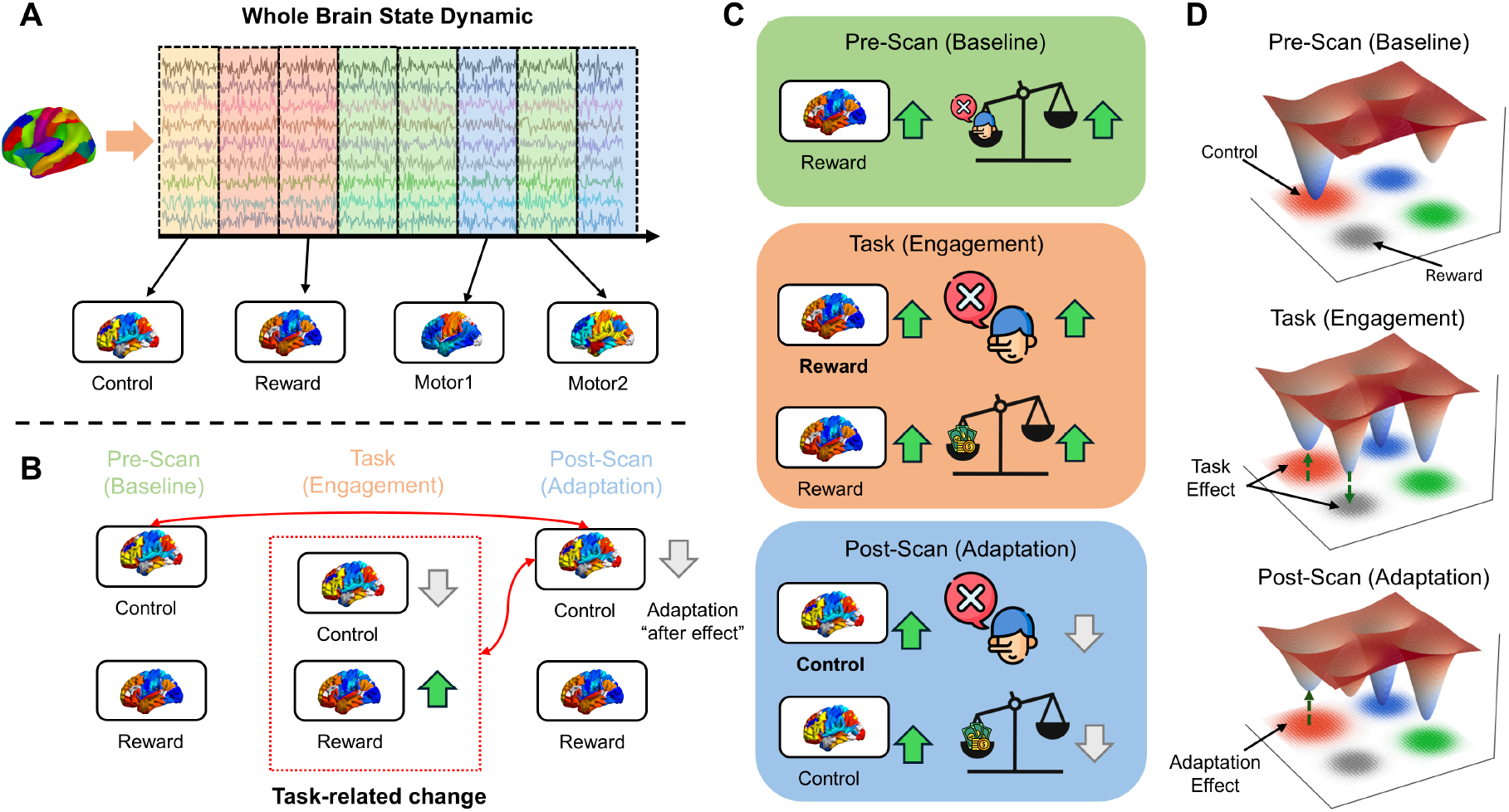
Graphical Summary: (A) In this study, we used HMM to extract 4 brain states that recur throughout the fMRI scan sessions. We label each HMM state as a control network, a reward network, and motor networks. Through Viterbi Pathways, the fMRI signal at each time point is labelled as one of the four HMM states. (B) The dynamic changes in the control network and the reward network, before, during, and after the task. Task-related changes include sustained activation of the reward network and inhibition of the control network. The task-induced after-effect is a persistent inhibition of the control network, resulting in a significant decrease in the proportion of control states after the task compared to the pre-task baseline. (C) The ability of brain states to predict behavior at different scanning stages. The higher the activation of the reward network in the baseline period before the task, the higher the bias towards dishonesty during the task. In the engagement period during the task, the higher the activation of the reward network, the higher the lying rate and the weighting of the reward. In the adaptation period after the task, the higher the residual control network, the lower the lying rate during the task and the lower the weighting of the reward. (D) The “attractor” model of moral brain states in moral dependence. Brain states are represented as attractors, and the attractor space is reconfigured at different stages of the moral task. During the task, the control network attractor becomes flatter, while the reward network attractor becomes more tilted, making it easier for states to stay in the reward attractor for longer periods of time. After the task, as the control network continues to be suppressed, the response attractor continues to become flatter as an adaptation effect.

Third, as dishonesty persisted, FO of the control network gradually decreased, while FO of the reward network progressively increased. Finally, pre-task resting-state activity and early-task activation of the reward network, as well as post-task activation of the control network, reliably captured individual differences in motivated dishonesty. Collectively, these findings provide new insight into how task engagement in dishonesty influence brain dynamics.

### 3.1 The moral disinhibition: dishonesty suppresses the control network and enhances the reward network

Previous studies have consistently shown that monetary rewards can increase the likelihood of dishonest behavior [40, 58]. In line with these findings, we find that different reward contingencies systematically influence levels of dishonesty. At both the session and condition levels, the enhanced lie condition significantly promotes dishonest behavior, exceeding the lie rate observed under enhanced honest and random conditions. This confirms that our information-sending task successfully elicited dishonesty.

Prior work have suggested that dishonesty can impact neural processes [18, 14]. Indeed, we observe evidence of brain state reconfiguration. As shown in the summary graph Figure 10B, such reconfiguration could be attributed to two aspects, the task-related changes and the adaptational after effects. During the task, our directional network-based analyses reveal that the transitions are predominantly directed toward the control network. Given that the task involved moral conflict, this reconfiguration toward the control network highlights the central role of cognitive control in managing internal conflict, suppressing automatic moral responses, self-regulation, and inhibiting unethical behavior [20, 41, 5]. As the task continued, the FO of the control network progressively decreased, while the FO of the reward network progressively increased. Unlike the control network, the reward system encodes the subjective value of outcomes and serves as a driving force for unethical behavior [24, 76, 19]. Therefore, this trajectory reflects the neural mechanisms underlying the escalation of dishonest behavior: With sustained dishonesty, the self-control resources gradually depleted, and individuals were unable to resist the monetary temptation and adapt to dishonest behavior, resulting in reward-related coding of decision outcomes becoming more pronounced. This task-specific brain reconfiguration is common and has been reported in previous studies, for example, when listening to different texts [82], in bilinguals [78], during learning[28], and in cognitive control [65]. The common interpretation is that this allows a task-specific processing and reflects the effort to allocate resources in response to external stimuli. In the context of dishonesty, this suggests that the dynamic interplay between control network and reward network reflects the ‘moral slippage’ effect, with a growing responsiveness to incentives for dishonest behavior, and diminishing capacity for self-regulation.

For the adaptational after effects, we found that the FO of the control network decreased after the task, and the overall switching rate among brain states was also reduced. This suggests that post-task brain dynamics were less inclined to engage the control network compared to baseline dynamics. When we consider the post-task brain dynamics as a window to examine the impact of the task on resting-state brain activity, this can be interpreted as reflecting the carry-over of the task-induced cognitive state.

Since the control network is suppressed as individuals adapt to dishonesty, this leaves a residual effect extended to the post-task session. Similar observations have been made in healthy individuals and clinical populations [38, 54]. For example, Roshchupkina et al. found that changes in the sensorimotor network during the post-motor learning period compared to the pre-task period reflect individual differences in motor learning [54]. Here, rather than returning to the baseline level, the control network remains suppressed after the task, suggesting that repeated involvement in dishonesty during the task leaves lasting traces in the dynamics of the brain. This sustained suppression may reflect lingering adaptations of the cognitive control system, which becomes less available for post-task regulatory processes. In this sense, the post-task control network functions as a neural marker of how dishonesty could have an “aftereffect” on neural dynamics beyond the immediate context of decision-making. Additionally, the observed reduction in switching rates indicates a stabilization of brain dynamics, potentially reflecting an efficient neural adaptation to task demands. Similarly, stabilization effects have been reported in other cognitively demanding tasks, where dynamic reconfigurations support optimized resource allocation [11].

### 3.2 Dynamic reorganization reflects individual differences in dishonesty

As expected, we find that the observed reconfiguration of brain dynamics is associated with individual differences in dishonest behavior. Our analyses show that the FO, dwell time, cumulative transitions, and interval time of the post-task control network are significantly associated with levels of honesty during the task. Specifically, stronger overall engagement with the control network, longer staying times per visit, and higher cumulative transition probabilities to the control network predict lower levels of dishonesty during the task. In contrast, less frequent visits (longer interval times) to the control network predict higher levels of dishonesty. These results align with the interpretation that the post-task control network serves as a neural trace of dishonesty-related dynamics [20, 41, 5]. In other words, individuals who continued to engage the control network even after the task were those who resisted dishonesty more during the task, which makes them less adapted to dishonesty. Whereas those who disengaged from the control network more strongly during the post-task period are those who succumbed to dishonesty. Previous post-task fMRI studies have also shown similar results consistent with our findings. For example, Tung et al. found that motor tasks modulated the resting-state brain, with the cross-correlation coefficient between the left and right motor cortices in the post-task resting scan being higher than before the task ([70]). These findings also held true for large-scale resting-state brain networks.

Pyka et al. found that the difficulty of a working memory task modulated post-scan DMN activation ([50]). Furthermore, behaviorally, studies have shown that different lingering mnemonic biases can be induced depending on the previous task, such as recent encoding of novel objects or retrieval of old objects ([15]). Our study extends these findings to brain state, highlighting the lingering effect between post-task resting-state brain state and both task and behavior.

The complimentary drift diffusion modeling(DDM) analysis quantifies participants’ weighting of monetary rewards and bias toward decisions. This reveals the relationship between brain state and behavior before and during a task. We find that in the early task phase, higher FO, dwell time, and cumulative transitions to the reward network are related to stronger weighting of monetary value. Likewise, higher cumulative transitions to the reward network during pre-task resting state were related to greater bias toward dishonesty across sessions. This supports our interpretation of State 1 as the reward network. Previous studies have demonstrated that the strength of reward-related activity predicts dishonest tendencies, reflecting the increased sensitivity to monetary incentives [46, 1]. Furthermore, previous research has also pointed to a close relationship between value representation and motivated dishonesty [19, 24, 36]. Liang et al used fNIRS to find that individuals who are more sensitive to rewards are more likely to lie [36]. Accordingly, stronger reward processing is associated with greater motivation to behave dishonestly. Finally, we also found that post-task interval times of the control network are linked to monetary weighting: individuals who prioritized monetary rewards during the task subsequently showed less frequent engagement of the control network (higher interval times) in the post-task period.

Interestingly, these associations are present only in Session 1 and Session 4. That is, during the initial and final stages of the moral task. One possible interpretation is that, early in the task, participants are still calibrating their behavioral strategies such that neural dynamics more strongly capture their inherent tendencies toward honesty or dishonesty. By the final stage, cumulative effects of dishonesty, such as rationalization and habituation, reveal individual differences in how the control and reward systems adapt [69, 27]. In contrast, mid-task sessions may reflect transient periods of adjustment, where neural dynamics are more unstable and therefore less predictive of individual differences [30, 85].

As shown in Figure 10C, we summarize the relationship between the brain dynamics and immoral decision as follows: The pre-task scan reflects a cognitive baseline, the task scan captures changes induced by task engagement, and the post-task scan indexes subsequent adaptation to the task [35, 75, 48, 8]. At baseline, responsivity of the reward network predicts individual propensity to lie during the task, highlighting a relationship between dishonest individual differences and baseline sensitivity to reward. During task engagement, activation in the reward network is associated with lying rates and the weighting of monetary incentives, emphasizing that repeated dishonest behavior is governed by reward-driven decision processes. Critically, this adaptation extended beyond the task itself, manifesting as persistent alterations in post-task control network dynamics. The control network predicts honesty rates and the weighting of reward representations, suggesting that it plays a central role in suppressing dishonest responses and modulating cognitive processing that prioritizes reward. Overall, the dynamic interactions between the reward and control networks jointly reshape individuals’ decision policies and behavioral tendencies when confronted with incentive conflicts.

### 3.3 How network dynamics changes: a Moral-dependent attractor model

By characterizing pre-task, task-engagement, and post-task brain states, we propose in Figure 10D a moral-dependent attractor model of the brain states. According to a current theoretical perspective, brain responses can be mapped onto an energy landscape, where the lowest energy point corresponds to stable, self-sustaining neural activity patterns that instantiate distinct brain states [37, 29, 43, 61]. Crucially, recent work demonstrates that the geometry of the attractor landscape undergoes systematic changes in response to external task demands [61]. Therefore, the brain dynamics identified can be represented by attractors. During the pre-scan baseline, the control attractor occupies a deep valley in the energy landscape, reflecting a stable and readily recruitable control state. During task engagement, task demands flatten the control attractor—reducing its depth and making neural trajectories less likely to converge onto the control state—while steepening the reward attractor, which increases the likelihood that neural activity will be captured by reward-related dynamics. Under post-task adaptation, the control attractor remains in the shallow region of the landscape, reflecting the depletion of control resources [41, 5]. This account provides evidence for context-dependent re-prioritization of neural states and highlights a dynamical vulnerability whereby reward-driven attractors can bias the system away from honest (control-driven) responses. In sum, the moral-dependent attractor framework explains how large-scale brain activity reconfigures throughout different stages involving moral conflicts.

### 3.4 Limitations and future perspectives

While our findings contribute to existing literature by comparing pre-task, during-task, and post-task dynamics, several limitations should be acknowledged. First, HMM is a statistical technique rather than a biophysical model of neural activity. The model assumes discrete brain states rather than continuous mixtures. Furthermore, the four-state solution may not represent the optimal decomposition across all datasets and populations, as the parameter *K* was selected through a data-driven approach that balances stability and representativeness of brain activation patterns. Indeed, there are other HMM solutions in the literature, including a 3-state solution [37], a 6-state solution [45], a 10-state solution [42], a 12-state solution [73], and a 19-state solution [66]. Future research should therefore replicate HMM analysis across multiple datasets and scanning sessions to explore the generalizability of HMM-derived dynamics. Lastly, the present study is more focused on dishonesty motivated by monetary reward. We have not investigated other motivational factors that drive immoral behavior. Future work should explore whether different motivational factors involve similar or different neural dynamics in short- and long-term periods.

## 4 Conclusions

In conclusion, this study demonstrates that task engagement in immoral decisions significantly reshapes resting brain dynamics. By applying a four-state HMM to fMRI data, we show that dishonesty is associated with reduced engagement of control networks, enhanced engagement of reward networks, and long-lasting alterations that extend beyond the task itself. Moreover, these dynamic reconfigurations not only track behavioral dishonesty but also capture individual differences in sensitivity to monetary rewards and bias tendency toward dishonesty. Together, our findings advance understanding of the adaptive neural mechanisms through which dishonesty reshapes cognitive control and reward processes in immoral decision-making.

## 5 Methods

### 5.1 Participants

Thirty-seven healthy participants (31 females, mean age = 23.27 ± 3.52 years) were recruited at the University of Macau. Participants were recruited from the student community at the University. All participants were right-handed and had no history of neurological or psychiatric disorders. Participants gave written informed consent after a complete description of the study was provided. Participants received a full debriefing upon completion of the experiment. All procedures involved were in accordance with the Declaration of Helsinki and were approved by the Institutional Review Board (IRB) of the University of Macau (BSERE21-APP005-ICI). The sample size for the fMRI study was determined based on previous fMRI studies on dishonesty (e.g., *n* = 28 [18], *n* = 40 [64], *n* = 25 [67], *n* = 31 [47]).

### 5.2 Task design

The information-sending task (IST) used in this study was part of a series of experiments spanning nine sessions[79] (see details in supplementary materials). From the participants’ perspective, the entire experiment was focused on receiving and passing information. The focus of our study was on the restingstate fMRI (rs-fMRI) before and after the main information-sending task. As shown in Figure 1, during IST, participants were asked to pass the food preference information of Mr. Li (36 pairs of food) to the next participant (the information receiver) with the consideration of reward units in four scan sessions. We manipulated the reward unit differences between two items, yielding three conditions: enhanced dishonesty, enhanced honesty, and random. Under enhanced dishonesty and honesty conditions, rewards were consistently higher for the correct or the error choices, while under the random condition, rewards were randomly designed. Each condition comprised 8 repetitions for each item, with 2 repetitions for each run. The enhanced honest, dishonest, and random conditions consisted of 72 trials separated into four runs.

### 5.3 fMRI Image collection

All images were acquired on a 3T Siemens Tim Trio scanner with the 12-channel head coil. Functional images employed a gradient-echo echo-planar imaging (EPI) sequence with the following MRI scanner parameters: (TE = 40*ms*, TR = 2*s*, flip = 90^*°*^, FOV = 210*mm*, 128 by 128 matrix, 25 contiguous 5*mm* slices parallel to the hippocampus and interleaved). We also acquired the whole-brain T1-weighed anatomical reference images from all participants (TE = 2.15*ms*, TR = 1.9*s*, flip = 9^*°*^, FOV = 256*mm*, 176 sagittal slices, slice thickness = 1*mm*, perpendicular to the anterior-posterior commissure line).

### 5.4 fMRI Imaging preprocessing

Image preprocessing was carried out using a standard pipeline called fMRIPrep(https://fmriprep.org/) [16]. Structural images were first corrected for intensity non-uniformity caused by magnetic field inhomogeneity using N4BiasFieldCorrection and spatially normalized to MNI152NLin6cAsym space. Brain tissues, including grey matter, white matter, and cerebrospinal fluid (CSF), were also segmented. For the functional images, motion correction and slice-timing correction were applied, and then they were aligned to the structural image. For denoising, we followed the same pipeline as listed in [80]. A confound regressor was built with 18 confound parameters using Python’s Nilearn library, including 6 motion parameters (*trans*_*x*_, *trans*_*y*_, *trans*_*z*_, *rot*_*x*_, *rot*_*y*_, *rot*_*z*_), global signal, framewise displacement, six anatomical CompCor confounds, and four discrete cosine-basis regressors. Temporal preprocessing included filtering the data with a low-pass filter of 0.08 Hz and a high-pass filter of 0.009 Hz to remove high-frequency noise and noise associated with scanner instabilities.

### 5.5 HMM analysis

HMM analysis was performed on two sessions of rs-fMRI data and 4 sessions of task fMRI data using the HMM-MAR toolbox developed by OHBA (https://github.com/OHBA-analysis/HMM-MAR). The HMM takes a set of multidimensional input time series and models the observed time series into a collection of hidden states. These hidden states recur over time according to a time-invariant state transition probability matrix. Specifically, the HMM consists of two parts: a set of hidden states (*k*) whose dynamics are governed by a transition probability matrix, and an observation mode, which is the process of generating data given the hidden state. In the present study, the observation model was defined as a multivariate Gaussian, such that the probability distribution of the observed data given a particular hidden state follows a normal distribution.

For the input features, we employed a 100-region parcellation of the 7 canonical networks (visual, somatomotor, dorsal attention, ventral attention, limbic, frontoparietal, and default) derived by Schaefer et al. [55]. The rationale for using this parcellation scheme is twofold. On the one hand, previous research indicates that overly fine-grained parcellations would result in high model complexity, as estimating potentially billions of parameters would far exceed the available data points [2]. On the other hand, coarse parcellations at the network level may fail to capture the temporal dynamics, since substantial temporal variance may occur within the large-scale networks. Thus, the 100-parcel resolution strikes a balance between model fitting and sensitivity to meaningful dynamic fluctuations [66].

Variational Bayes (VB) was employed to obtain an approximation of model posterior distributions. VB infers the model parameters by alternating a variational expectation step and maximization step. Specifically, it randomly initializes an approximate distribution for model parameters and calculates the variational free energy as a cost function, which contains three terms: the averaged log-likelihood, the Kullback– Leibler divergence, and negative entropy. The aim here is to minimize the cost function. With reference to the previously established procedure, we chose to set the training cycle to 1000, and we repeated inference with 300 cycles with different initializations [62, 75]. The cycle with the lowest free energy was chosen as the starting point of HMM inference [73, 42, 62]. After comparing a range of ***K*** values in terms of robustness and degree of fit to the data, we selected the model solution with 4 hidden states (*see supplementary materials for more details*).

Then, the estimated transition and the observation model were applied to obtain the most likely sequence of hidden states using Viterbi algorithm. It estimates the probability of each latent state being the most likely state at a specific time point. Here, we chose the state with the highest probability and defined that given time point to belong to such a state. From here, we summarized the characteristics of each subject-specific HMM in terms of FO (the proportion of time each participant spent in each brain state), dwell time (the average duration in one visits to a certain state) and interval time (the average duration between visits to the same state), Switching Rate (frequency with which participants transitioned between all brain states) and State Transition Probability.

### 5.6 Neurosynth Decoding

To quantify the functional relevance of the inferred hidden states, we used the Neurosynth database, which contains nearly 14,300 fMRI studies and 507,000 reports of BOLD-inferred brain activity [83]. A decoding model was trained on the database that maps keywords extracted from the body of literature to the locations of brain activity. Here, we performed forward association decoding *P* (*Activation*|*State*) to map the spatial expression of each state onto 13 Neurosynth topics, which encompass a variety of brain processes applicable to our task.

### 5.7 Statistical comparison and correlation analyses with HMM dynamic

The comparison between pre, during and post-task HMM dynamic metrics was assessed using paired-sampled Wilcoxon signed-rank. For examining the HMM dynamics within the 4-task session, repeated ANOVA was used. For linking HMM states with behavior, Spearman’s rank correlation analyses was used. All p-values were corrected for multiple comparisons using the false discovery rate unless otherwise specified.

### 5.8 Drift Diffusion Modeling

We used a Bayesian hierarchical drift-diffusion model (HDDM) framework suited for estimating trial-by-trial parametric modulations on latent decision processes[77]. Following a standard procedure of HDDM model estimation, we used Markov chain Monte Carlo (MCMC) sampling methods for Bayesian approximation of the posterior distribution of parameters (generating 100,000 samples, discarding 5000 samples as burn-in[53]). Here, dishonesty responses were coded as 1 and honesty choices were coded as 0. Reaction times (RT) longer than 20 s or shorter than 0.3 s were excluded[86]. We inspected traces of model parameters, their autocorrelation and computed the R-hat (Gelman-Rubin) convergence statistics to ensure that the models had properly converged[77]. Five chains were run, each with 11000 iterations and 1000 burn-in samples. No R-hat statistics were larger than 1.1, indicating good convergence[71]. Parameter distributions at both the group level and the individual-participant level were simultaneously estimated[72]. Deviance information criterion (DIC), suitable for hierarchical model comparison, was used as a measure of goodness-of-fit[77]. To evaluate if the preferred model can reproduce key patterns in the observation, we carried out posterior predictive checks, where we simulated data based on the parameters derived from the preferred model. This analysis showed that the preferred model satisfactorily reproduced the observed proportion of dishonesty choices and the means and the quantiles of RT (supplementary table 1). This indicated that the preferred model could reliably reconstruct the patterns in the observed data.

## Acknowledgements

This work was mainly supported by the Science and Technology Development Fund(FDCT) of Macau [0112/2024/RIA2], and MYRG of University of Macau(MYRG-GRG2025-00175-ICI). The authors thank all research assistants who provided general support in participant recruiting and data collection.

## Ethical Statement

The authors are responsible for all aspects of the work in ensuring that questions related to the accuracy or integrity of any part of the work are appropriately investigated and resolved. All procedures performed in this study involving human participants were in accordance with the Declaration of Helsinki and were approved by the Institutional Review Board (IRB) of University of Macau (BSERE21-APP005-ICI).

## Data and Code availability

The data is available upon request from the corresponding author. The data analysis code is available at

https://github.com/andlab-um/uaHMM.

## Notes

### Competing Interest Statement

The authors have declared no competing interest.

### Summary of Updates

This version of the manuscript has been revised to update the changes in the introduction and discussion to reflect more on the impact of task engagement on brain dynamics. Figure 1A has been revised slightly to 2 hypotheses instead of 4

