## Supplementary files for "Task engagement in immoral behavior altered post-task brain state"

The alternated brain states in resting state after immoral decisions

**This PDF file includes:**

Supporting text

Figures S1 to S11

Table S1

SI References

Supporting Information for Task Instructions:

Welcome session Welcome! This is an experiment about information receiving and passing decisions. We are interested in understanding how people make decisions about receiving information from others and how people send information to others. There are a lot of people participating in this experiment, and everyone is required to receive and pass information about a stranger. It should take about an hour and 45 minutes, including instructions, testing. For participating, you will receive at least ¥80. Depending on your choices during the task, you will have the opportunity to earn from ¥0 up to ¥100 more. You will be paid in cash for your time and your earnings at the end of the experiment.

Please select one envelope here and tell the number of the envelope you choose without opening it. After this, we will go to the first step of the study.

**Step 1: The first food preference task**

In this task, there is no such thing as a correct or incorrect answer, and your choices won’t have any consequences (reward or punishment). This task is about choosing the food you prefer between two choices and then rating your preference from 1 to 100. The task will be performed using the mouse. First, you need to press the ‘Start’ button, and two foods will appear in the upper left and right corners. Please respond within 4 seconds, or you will receive a time-out warning. After clicking the preferable food, you need to rate your preference level from 0 to 100. If you are ready, we can start now.

**Step 2: The first information-receiving task with mouse tracking**

This step is a part of information sending, where you receive information about a stranger from the last participant. The format is similar to the first step. First, the information category will be shown in the center of the screen (e.g., name). Then you press the ‘Start’ button, and two pieces of information will appear in the upper left and right corners. The information sent from the last participant will be asterisked, which could be correct or incorrect. You need to choose whether to trust this information or not by clicking one of the choices, since you know nothing about the stranger. Please respond in 4 seconds, or you will receive a time-out warning. If you choose the wrong answer, you will lose 1 point in this experiment. Your final earnings are positively correlated with your experiment score. If you are ready, we can start now.

**Step 3: The first food memory task with mouse tracking**

Please open the envelope and memorize the favorite foods of Mr. Li, which are very important for our task. The form of the memory test is similar to the food preference task, where you need to choose between two food pictures using the mouse. If you think you have already memorized Mr. Li’s favorite foods, let’s start the practice session. In the practice session, you will receive feedback if you choose the wrong answer. Now is the formal test session without feedback. In this test, your mouse movement will also be recorded, so please respond within 4 seconds, or you will receive a time-out warning. After choosing the answer, you need to rate your confidence in your choice. You should rate your confidence seriously, even though it doesn’t have any consequences. Remember that we will move to the next step only if your accuracy reaches 90%. If you are ready, we can start now.

**Step 4: First Resting fMRI Scan**

In this step, we will conduct our first resting fMRI scan. Please close your eyes, relax, remain as still as possible and please do not fall asleep. The scan will last approximately 8 minutes. Are you ready? If so, we'll begin.

**Step 5: The information sending task (IST)**

In this step, you are the information sender, where you need to pass the food preference of Mr. Li to the next participant. On every trial, foods will appear on the left and right upper corners. Under each food, there is a bar indicating the reward (1 point to 9 points) you will receive if you choose this food. The next participant will receive the information you send about Mr. Li and perform a test similar to what you did in step 2. Generally, most people tend to trust you when they know nothing about Mr. Li. Their choices will affect their final earnings, and your reward in this step will affect your final earnings as well. This task consists of 4 blocks, meaning you need to pass the information 4 times. If you are ready, we can start now.

**Step 6: Second Resting fMRI Scan**

Same as step 4.

**Step 7: Exit questionnaire**

Congratulations! You have finished our task and thank you for your participation. Now please complete the questionnaire on the Wenjuanxing platform, which lasts around 30 minutes.

**Step 8: The surprise second memory test with mouse tracking**

This is a surprise session. We will test your memory about Mr. Li’s favorite food again. The procedure is the same as before, where you need to choose the correct answer and rate your confidence seriously. You should keep your accuracy as high as possible. If you are ready, we can start now.

**Step 9: The second receiving and trust decisions with mouse tracking**

Now we present the information sent from the last participant again to examine whether you still trust her/him or not. The procedure and requirement are the same as before, where you click the correct answer to gain more earnings, with the asterisked choice from the last participant. If you are ready, we can start now.

**Step 10: The second food preference task with mouse tracking**

Same as the first task.

**Step 11: Three days after the scanning: the third memory test**

Same as the surprise second memory test.

Within the scope of the current study, we focus our analysis on the 2 resting scans acquired in step 4 and 6, as well as the task scans acquired in step 5.

Supporting Figures S1 to S11

Fig. S1.

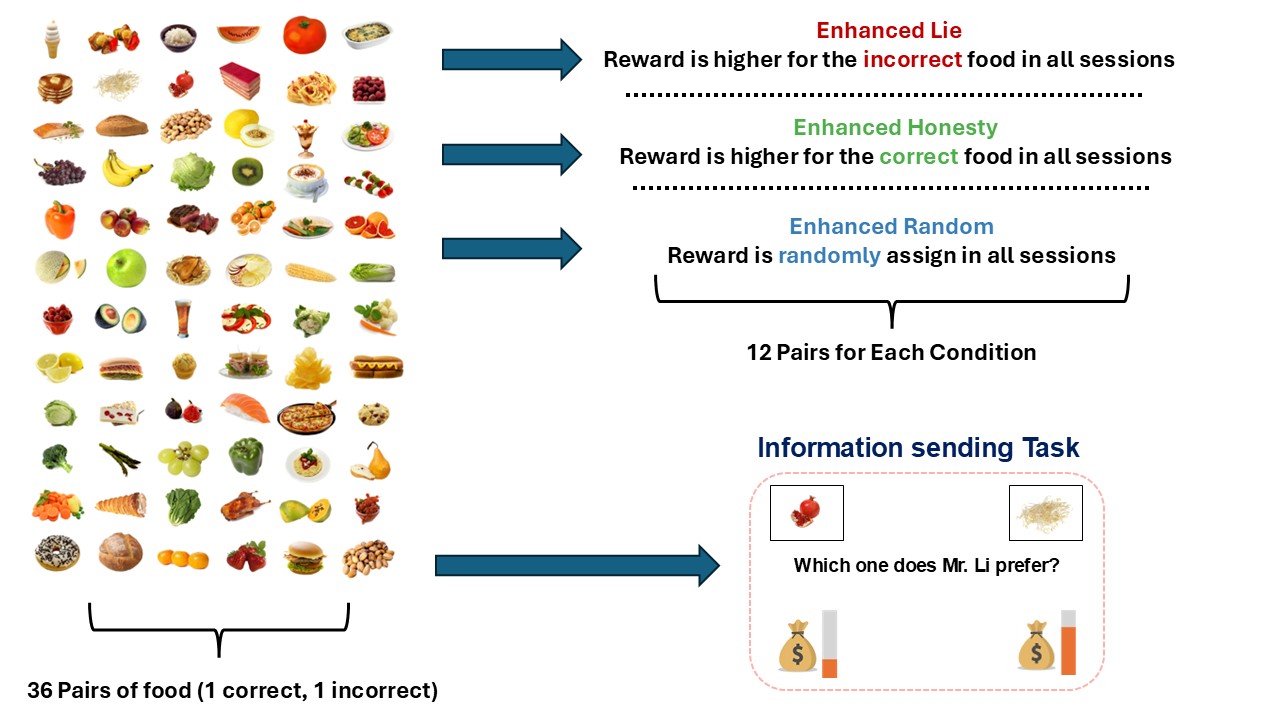
**Supplementary Figure 1**: Food stimuli used in the memory task are selected from “Food-Pics_Extended” (http://food-pics.sbg.ac.at). There were 36 pairs in total, and during each information sending trial, one of the pair would show up, where it contains 1 correct answer and 1 incorrect answer. There were 3 conditions in total, enhanced lie, where there is higher reward for the incorrect food in all sessions. Enhanced honesty, where there is higher reward for the correct food in all sessions. And finally, enhance random, where reward is randomly assigned in all sessions. There were 12 pairs for each session

Fig. S2.

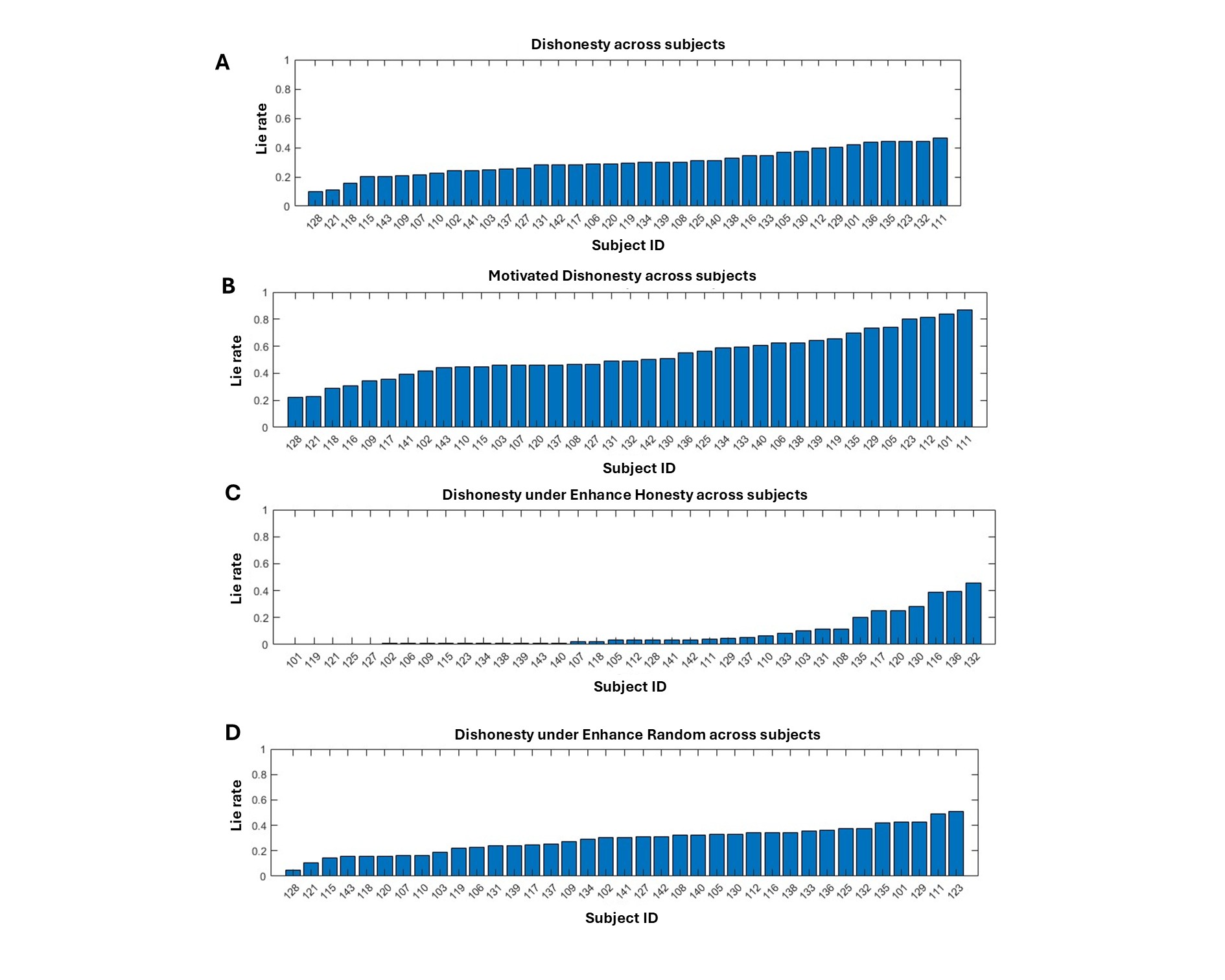
 **Supplementary Figure 2:** (A) Displaying the dishonesty rate across subjects averaged across all conditions. (B) Similarly, displaying the motivated dishonesty rate across subjects under the enhanced lie conditions. The lie rate varies significantly across 37 subjects. Some participants, even when faced with the temptation of rewards, exhibit an extremely low rate of dishonesty. In contrast, some almost choose to lie whenever the opportunity arises. This highlights the individual variation in motivated dishonesty in the face of reward. (C) Displaying the dishonesty rate under the enhance honesty condition (D) Displaying the dishonesty rate under the enhance random condition across subjects.

Fig. S3.

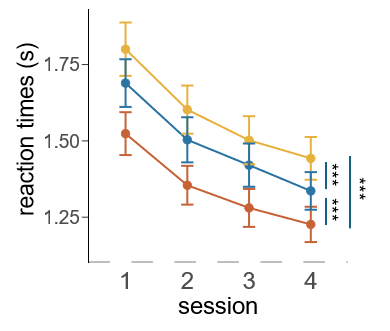

**Supplementary Figure 3:** Results of reaction time along the session in information sending task. *** denotes significant level at *p <* 0.001

Fig. S4.

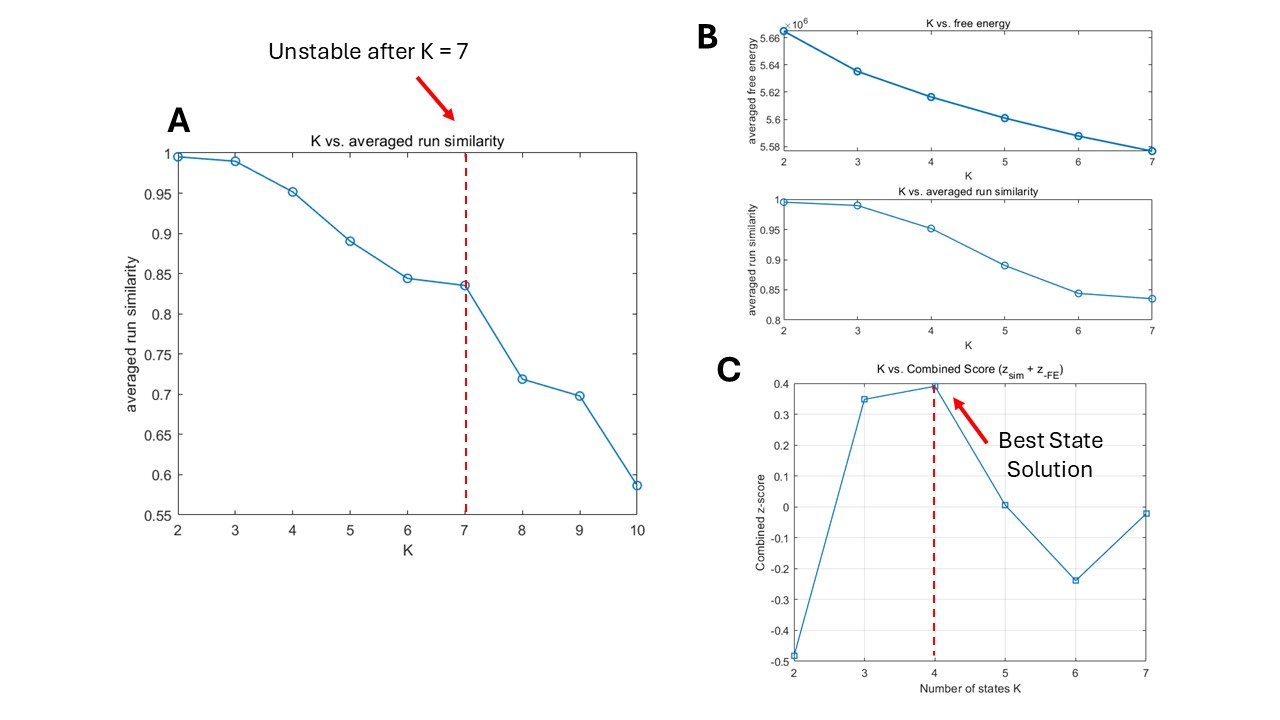

**Supplementary Figure 4**: (a) For each candidate K ranging from 2 to 10, we trained a Hidden Markov Modeling (HMM) model 5 times. With reference to previous research (van der Meer et al., 2020), we used the Hungarian method to estimate the pair-wise similarity between different repeats. Then, we took the averaged similarity as an indicator for the robustness of the estimated states (Li et al.,2024). We observe a general trend that as the number of state *K* increase, the model becomes more unstable, and the averaged similarity within a given HMM model eventually drop below 0.8 after *K* = 7. This is consistent with previous studies reporting that an increased K state value would lead to high model complexity, which results in capturing too much noise in the model (Li et al., 2024; Vidaurre et al., 2019). (b) Therefore, we limit our exploration between 2 state solutions to 7 state solutions. Theoretically, the choice of model should refer to model comparison metrics like free energy, however, previous research has shown that this metric tends to improve monotonically (Moretto et al., 2021; Van der Meer et al., 2020; liu et al., 2025). As expected, the free energy metric displayed here also shows to monotonically decrease. That is to say, as the model becomes more complex, the performance on the metrics becomes better, possibility because of overfitting. Thus, we refer to both free energy as well as the robustness of estimated states to choose the optimal state solutions for our data. (c) With reference to Liu et al (2024), we summed the z scored value of similarity and free energy. The result shows that when K = 4, the model strict a balance between model performance and robustness of state solution.

Fig. S5.

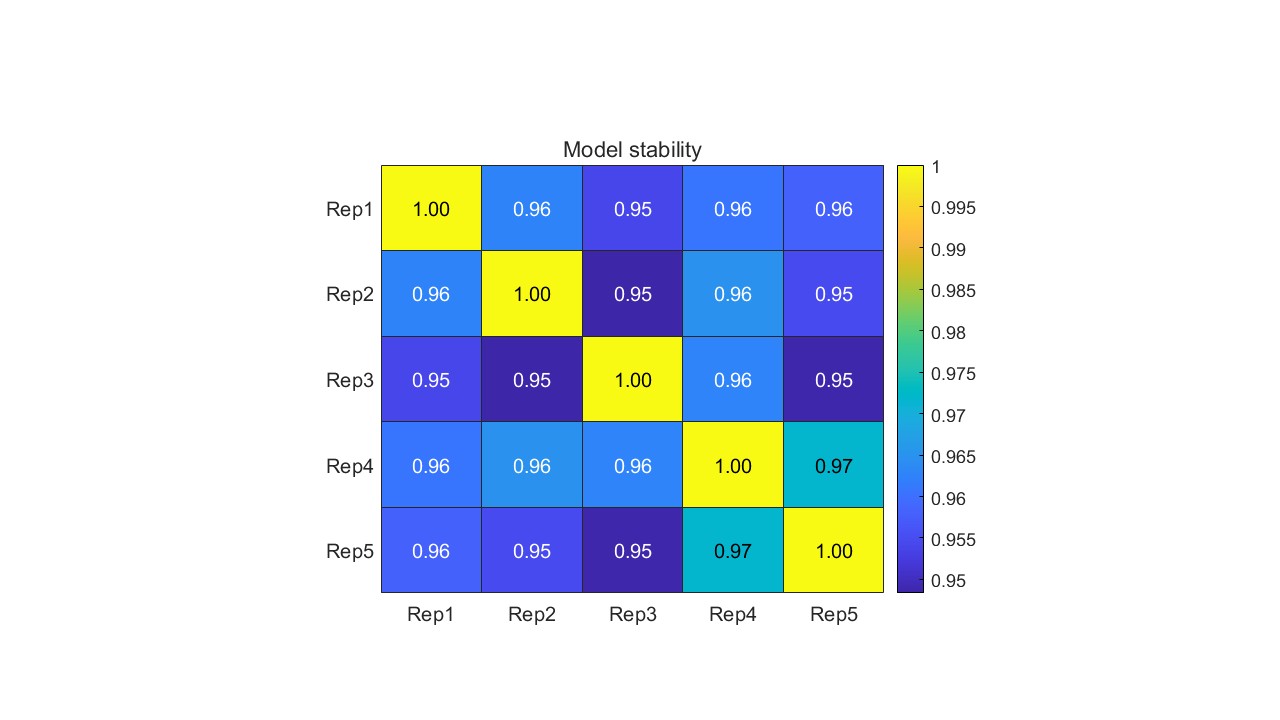

**Supplementary Figure 5**: To show the 4-state solution HMM is highly replicable and robust, we plot the pair-wise similarity between 5 repeated inferences for a 4-state solution HMM as model stability. As shown, the paired-wised similarity in 4-state solution HMM reached at least 0.95, suggesting that it was highly stable and robust.

Fig. S6.

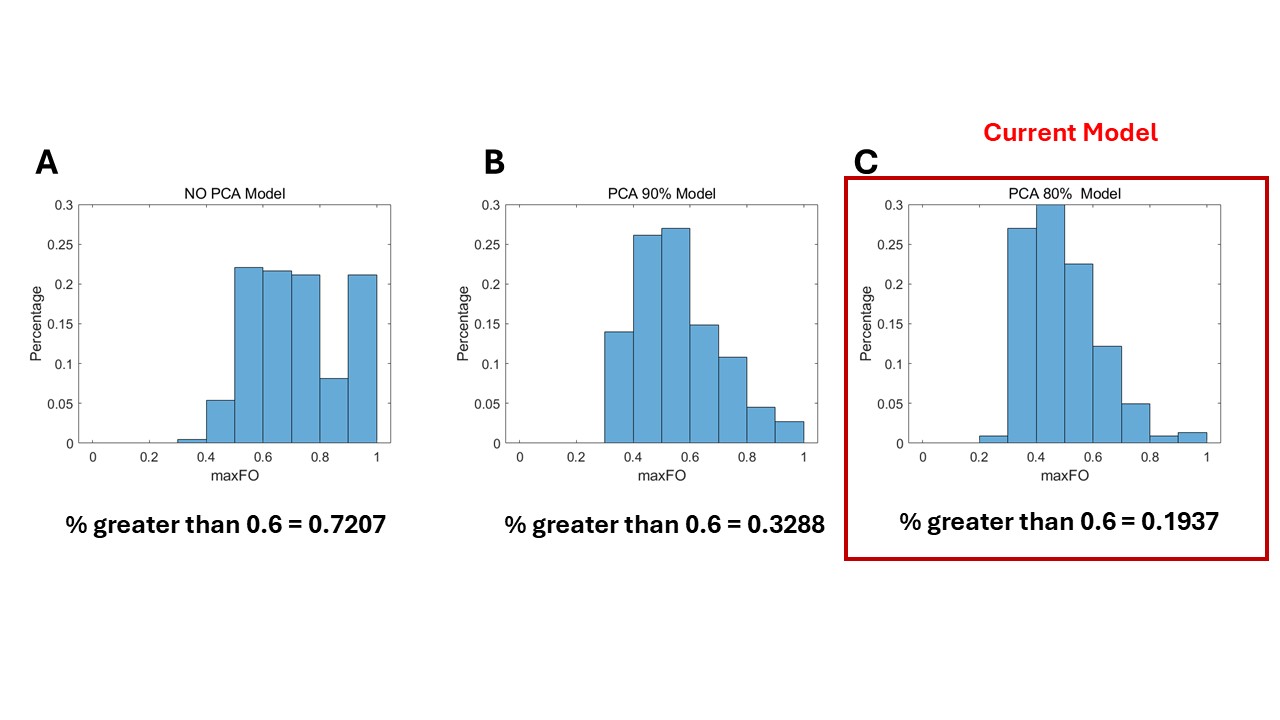

**Supplementary Figure 6**: (a) In addition, we also found that the HMM inference in the original fMRI space (Schaefer 100 atlas) would lead to serious model stasis. This was described in Ahrends et al (2022), that the model fails to capture any dynamic changes in the data, instead, the model assigns entire sessions to a single state, ignoring within-session dynamics. One metric describing the model stasis is maximum fractional occupancy (maxFO), this was defined as the maximum proportion that each state occupies in the time series of a particular subject. Therefore, the greater the maxFO, the more static the model is. As shown, models without PCA projection results in over 72% of the session to have a maxFO greater than 0.6. That is to say, most of the sessions are assigned to a signal state and the model fails to capture the brain state dynamics. (b) Therefore, we run all the HMM inferences in the PCA space, which has been shown by previous literature to reduce the static model (Zhang et al., 2024; Ahrends et al., 2022). Simply speaking, the main cause of model stasis was because there were potentially too many parameters to estimate from limited data. So reducing the dimension of input can significantly reduce the number of parameters that are required to be estimated, thereby reducing the model stasis. Specifically, we project the data from original fMRI space to a reduced PCA space and then conducted HMM inference within this reduced PCA space. Since PCA produces a projection matrix *P*, we can therefore reproject the PCA space back to the original fMRI space. However, models that work in PCA space that kept 90% of original variance still suffer from model stasis, namely, 33% of the session to have a maxFO greater than 0.6. Finally, we considered the model that operates in PCA space that kept 80% of the original variance, which shows a significant improvement. As shown, only 19% of the sessions had a maxFO greater than 0.6. Since further reducing the dimensionality could lead to excessive information loss, we conclude that maintaining an 80% PCA space for HMM inference strikes the optimal balance between preserving information and minimizing model stasis.

Fig. S7.

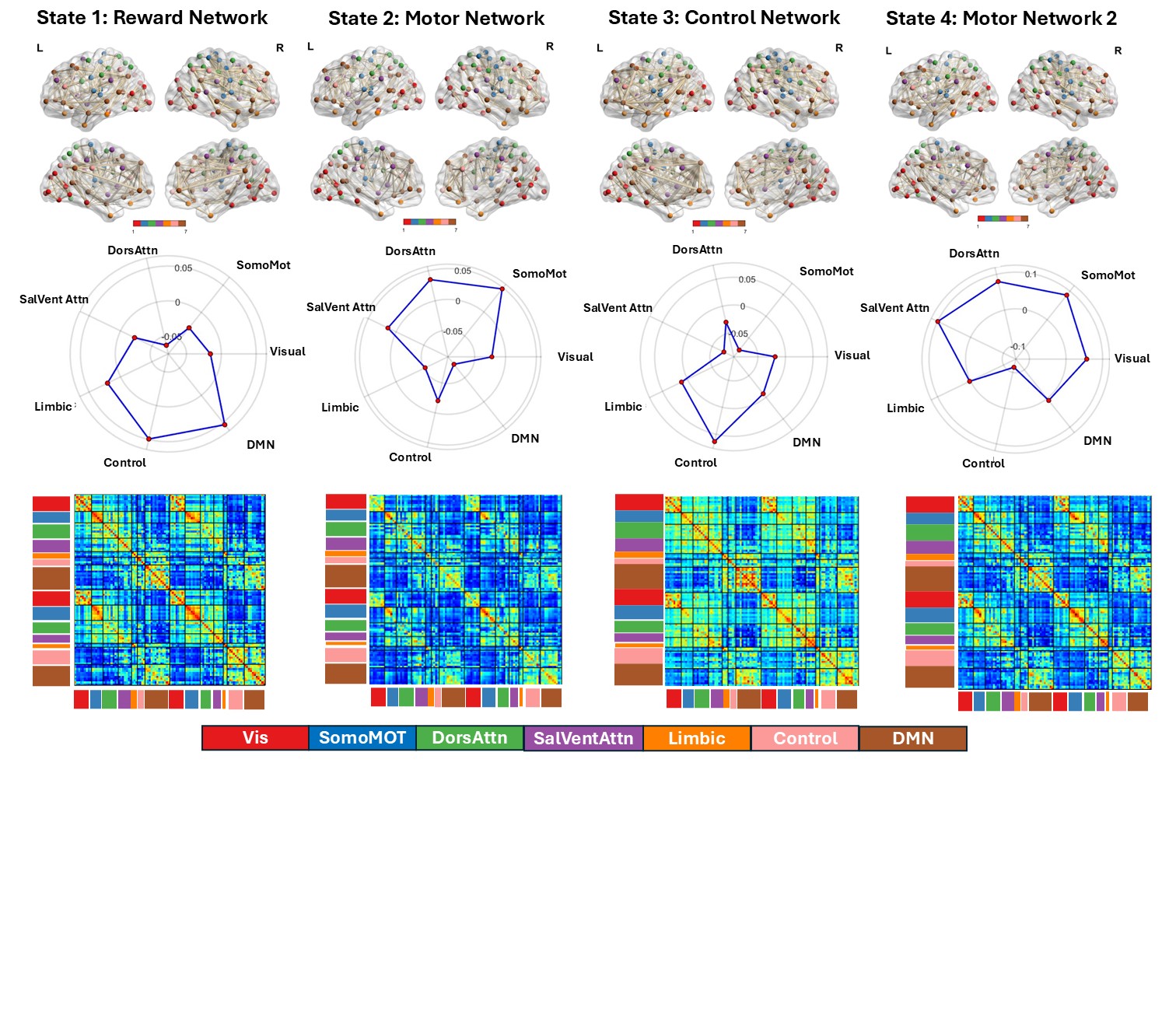

**Supplementary Figure 7:** Functional Connectivity and Activation on the 7 of the canonical brain networks in each of the 4 HMM states. The value here is obtained by the mean and covariance of the Gaussian distribution of the inferred brain state. Notes: Visual Network (Vis), Sensorimotor Network (SomoMOT), Dorsal Attention network (DorsAttn), Salience Ventral Attention Network (SalVentAttn), Limbic network (Limbic), Control Network (Control), Default Mode Network (DMN)

Fig. S8.

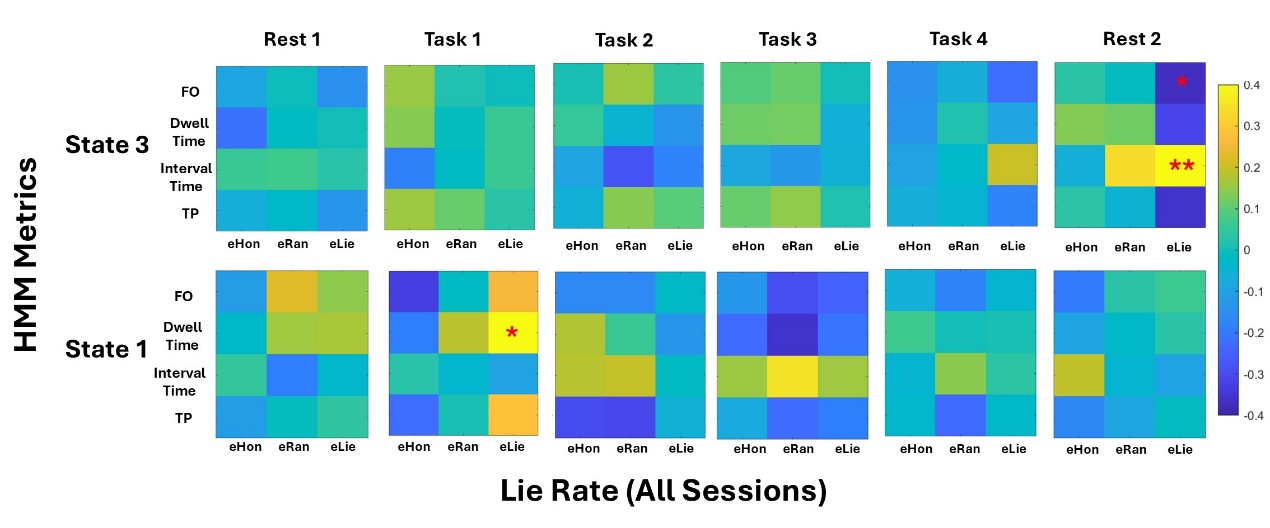

**Supplementary Figure 8:** Results of 4 HMM Metrics (FO: Fractional Occupancy, Dwell Time, Interval Time, TP: Cumulative Transitional Probability) across 6 session (Rest 1, Task 1, Task 2, Task 3, Task 4, Rest 2) and Lie rate averaged across all sessions in 3 conditions (eHon: Enhance Honesty, eRan: Enhance Random, eLie: Enhance Lie). Aster marks statistical significance after fdr correction: *0.05, **0.01

Fig. S9.

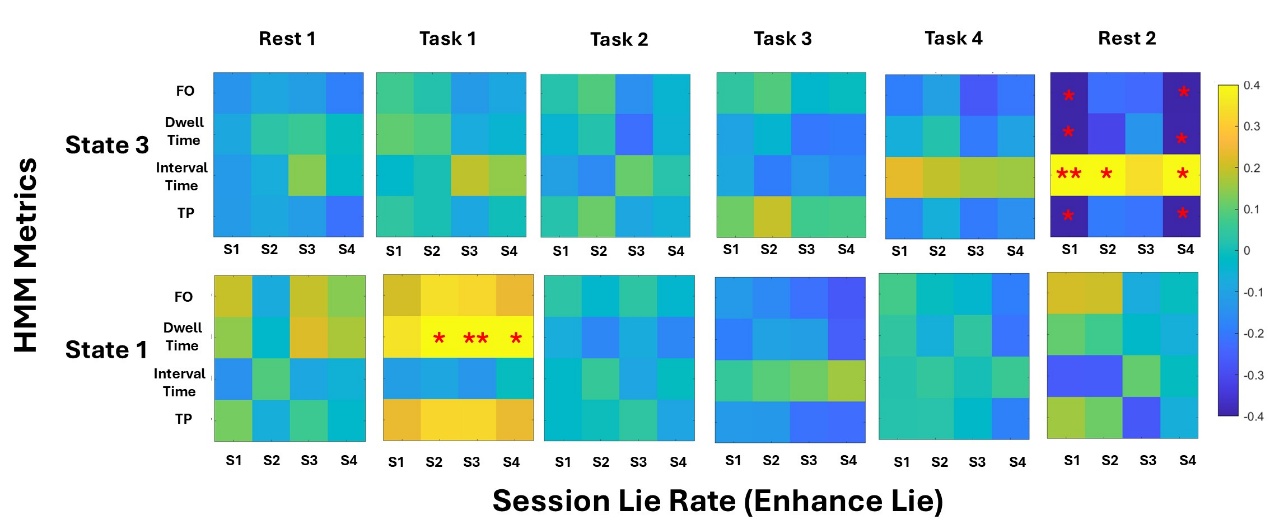

**Supplementary Figure 9:** Results of 4 HMM Metrics (FO: Fractional Occupancy, Dwell Time, Interval Time, TP: Cumulative Transitional Probability) across 6 session (Rest 1, Task 1, Task 2, Task 3, Task 4, Rest 2) and session lie rate in enhance lie condition. Aster marks statistical significance after fdr correction: *0.05, **0.01

Fig. S10.

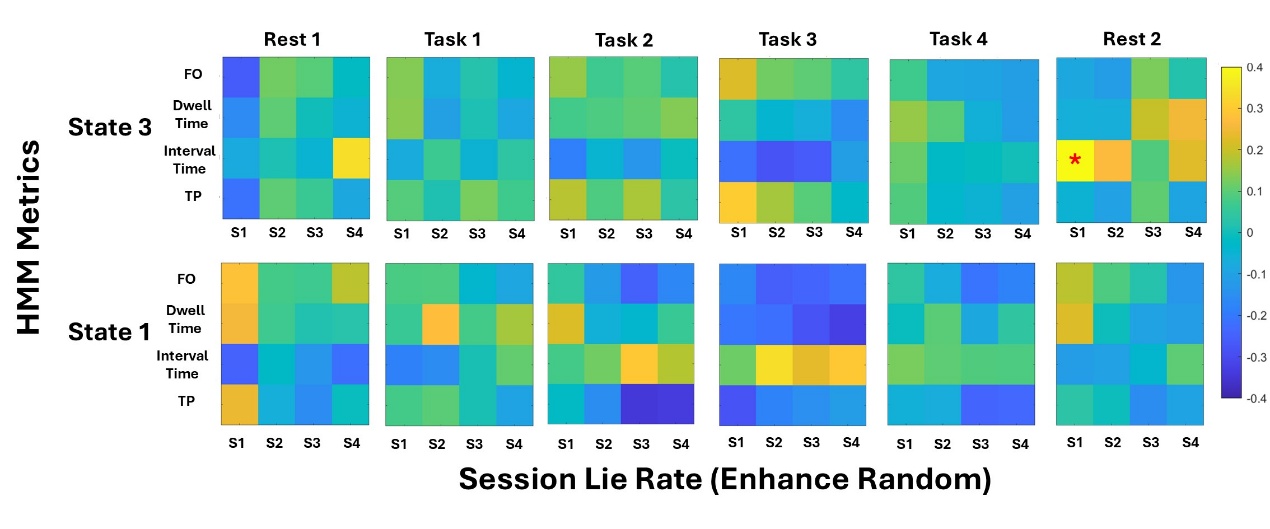

**Supplementary Figure 10:** Results of 4 HMM Metrics (FO: Fractional Occupancy, Dwell Time, Interval Time, TP: Cumulative Transitional Probability) across 6 session (Rest 1, Task 1, Task 2, Task 3, Task 4, Rest 2) and session lie rate in enhance random condition. Aster marks statistical significance after fdr correction: *0.05

Fig. S11.

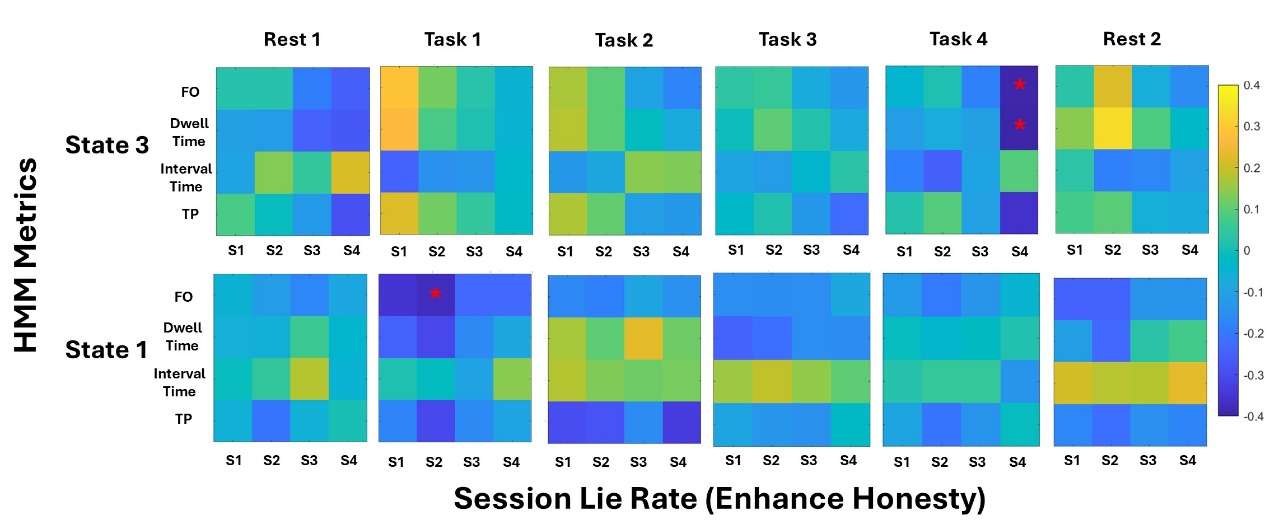

**Supplementary Figure 11:** Results of 4 HMM Metrics (FO: Fractional Occupancy, Dwell Time, Interval Time, TP: Cumulative Transitional Probability) across 6 session (Rest 1, Task 1, Task 2, Task 3, Task 4, Rest 2) and session lie rate in enhance honesty condition. Aster marks statistical significance after fdr correction: *0.05

Table. S1.

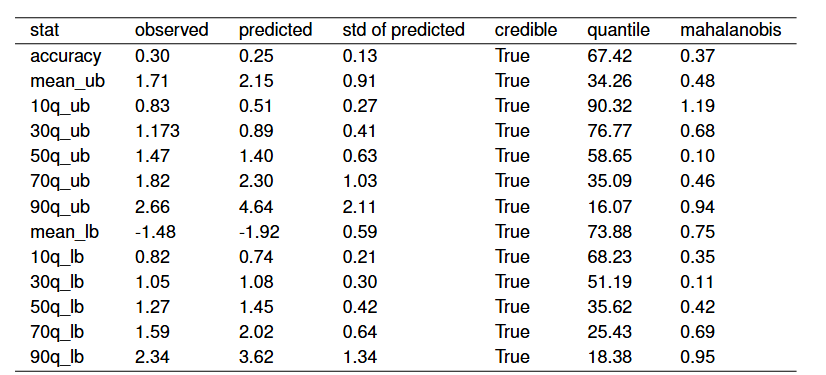

**Supplementary Table 1:** Posterior Predictive Checks for the preferred model (Model 5). Notes: lb: lower bound; ub: upperbound; 10q ~ 90q: 10th ~ 90th quantile of RT distribution; Credible: whether observed data falls in the 95% credible interval of the simulated data; Mahalanobis: Mahalanobis distance of the observed data from the center of distribution of the simulated data
